# Comparative responses of legume vs. non-legume tropical trees to biochar additions

**DOI:** 10.1101/2025.02.19.639164

**Authors:** Sean C. Thomas, Md. Abdul Halim, Syed Tuhin Ali

## Abstract

Nitrogen-fixing plants in the legume family (Fabaceae) may show particularly large positive responses to biochar additions due to their capacity to potentially compensate for reduced N in biochar-amended soils. Prior studies also suggest that biochar may have specific developmental effects on legumes, including increased root nodulation and altered morphology. We examined the growth and morphometric responses of legume and non-legume tropical trees to biochar additions in a common garden pot trial experiment. Four legume species (*Acacia auriculiformis*, *A. mangium*, *Delonix regia*, and *Pterocarpus santalinus*) and four non-legumes (*Eucalyptus alba*, *Melia azedarach, Swietenia macrophylla*, and *Syzygium cumini*) were compared in terms of sapling responses to additions of a wood-feedstock biochar applied at 10 and 20 t/ha. Overall, strong positive effects of biochar additions on sapling performance were observed, with an average increase of 30% in total biomass and a notable increase in height relative to diameter growth. Species showed pronounced differences in responses, with strong interactive effects of species and biochar treatments on growth metrics. Legume species showed an average increase somewhat greater than non-legumes; however, responses were variable among species, with the two *Acacia* species showing the largest responses, resulting in a non-significant pattern. A literature-based meta-analysis of tropical and subtropical trees likewise suggests greater biochar responses in legumes, but the analysis also falls short of statistical significance. In addition, experimental results indicate large interactive effects of species and biochar on soil pH and other soil properties. Large growth responses of certain taxa of legumes (and other taxa) to biochar, and pronounced species-specific effects on soil properties, may reflect evolved responses to fire disturbance that can be leveraged in the context of forest restoration and enhanced carbon sequestration in degraded tropical landscapes.

## 1. Introduction

Biochar, generally defined as pyrolyzed waste organic material designated for use as a soil amendment (Lehmann & Joseph, 2015), has recently received enormous research attention as a means to enhance carbon sequestration and improve the productivity of managed ecosystems, including forests. An important generalization to emerge on biochar use in managed ecosystems is that plant species vary greatly in their responses to biochar soil amendments. Such differences are evident in essentially all integrative analyses that have considered species-level effects, including meta-analyses of both agricultural crops (Jeffrey et al., 2011; Liu et al., 2013; Ye et al., 2020), and trees (Thomas & Gale, 2015; Juno & Ibáñez, 2021). By pooling data across many studies, meta-analyses potentially exaggerate apparent species differences in responses, since these differences will generally be confounded with differences in biochar feedstock and production parameters and dosages, as well as other experimental conditions. Nevertheless, published studies that have considered variability in species responses to the same biochar under the same experimental conditions have also documented high variability in species responses, including studies of early-successional herbaceous species (Gale et al., 2017) and trees (Sovu et al., 2012; Pluchon et al., 2014). Likewise, studies of biochar amendment effects on mixed-species vegetation have generally observed changes in species composition consistent with highly variable species-specific effects (van der Voorde et al., 2014; Bieser & Thomas, 2019; Williams & Thomas, 2023).

N-fixing plants, in particular legumes (family Fabaceae), might be expected to show particularly large increases in performance in response to biochar. During biochar pyrolysis most feedstock N is usually lost, and the remaining N is covalently bound and unavailable (Clough et al., 2013). In addition, biochar tends to strongly bind ammonium in the soil solution (Wang et al., 2015). Thus, biochar-amended soils initially show reduced N availability, though this may change through time as biochar alters N biogeochemical processes and enhances N retention in the system (Nguyen et al., 2017). N fixation is expected to alleviate reductions in the relative supply of N by providing N-fixing species access to the atmospheric N_2_ pool. In addition, the preponderance of evidence indicates that legumes generally respond to biochar by increasing root nodulation and N fixation (Farhangi-Abriz et al., 2022). This phenotypically plastic response is expected to further enhance the performance of N-fixing legume species in biochar-amended soils. Some studies have also noted large effects of biochar on other developmental patterns in legumes, such as a >100% increase in stem allocation and reduction in root allocation observed in *Leucaena leucocephala* in response to biochar additions (Thomas et al., 2019). However, pronounced changes in growth form have also been noted in non-legume trees; for example, Sifton et al. (2022) noted increases in the height:diameter ratio in *Acer saccharinum* in response to biochar.

Prior studies in the context of agriculture and ecological restoration suggest disproportionate responses of legumes to biochar. Large average growth responses in legumes were noted in the earliest biochar meta-analyses. Jeffrey et al., (2011) reported that soybean was one of only two crops that, considered individually, showed a positive effect of biochar on crop productivity. Liu et al., (2013), with an increased sample size of studies, found that among major crops, legumes showed an average 30% increase in productivity, compared to 7-11% for major cereal crops; however, at least one recent meta-analysis did not observe greater responses in legume crops (Farhangi-Abriz et al., 2021). Field studies of biochar effects on plant communities have likewise commonly noted disproportionate increases in cover or abundance of legume species (van der Voorde et al., 2014; Bieser & Thomas, 2019; Williams & Thomas, 2023). Studies of species mixtures have also found increases in non-legume species growth in response to the combination of N-fixing legumes plus biochar (Liu et al., 2017; Thomas et al., 2019; Sifton et al., 2022), attributed to inputs of soil N by legumes.

Responses of tropical trees to biochar are of particular importance and interest in several respects. Tropical forests play a major role in the global carbon cycle (Pan et al., 2011); however, most regions of the tropics have been subject to widespread deforestation and forest degradation such that carbon gain in secondary forests is critical to future tropical forest carbon sinks (Heinrich et al., 2023). Forest productivity, and thus the C sink, is often limited by soil nutrient status in degraded tropical landscapes (Powers & Marín-Spiotta, 2017). Biochar can thus potentially aid in forest restoration on degraded tropical soils, provided that trees show strong positive growth responses. Early studies suggested that tropical trees generally show large growth enhancements in response to biochar additions (as reviewed by Thomas and Gale 2015). However, results from recent studies, including field trials, are mixed, with studies providing evidence for both neutral (Gonzalez Sarango et al., 2021) and positive (Rios Guayasamín et al., 2024) effects on overall growth responses, depending in part on soil conditions. As with other ecological groups, species-specific differences in responses to biochar appear to be the rule in tropical trees (e.g., Sovu et al., 2012; Ghosh et al., 2015; Lefebvre et al., 2019; Rios Guayasamín et al., 2024). N-fixing legumes are an important component in tropical forests, particularly so in disturbed forests and in seasonal tropical and subtropical forest types (Gei et al., 2018; Tamme et al., 2021); however, studies to date have not addressed whether there is a systematic difference in biochar responses between legumes and non-legumes among tropical trees. Better information on species-specific responses of tropical trees to biochar, as well as potential systematic differences among phylogenetic and ecological groups, is of general importance in the context of tropical forest restoration and enhanced C sequestration.

In the present study, we assessed biochar responses of tropical trees on a representative degraded soil in the Sylhet region of Bangladesh, comparing responses of four legume and four non-legume tree species. We address the following questions: (1) Does biochar enhance overall sapling performance? (2) Do legume species show greater increases in biomass and other measures of plant performance than non-legumes? (3) Are there biochar effects on allometric or morphometric patterns and do legumes differ from non-legume species in this regard? (4) Are there positive effects of biochar on root nodule production in the legume species? (5) How do results compare with those of a meta-analysis of available data on tropical tree responses?

## 2. Material and methods

### 2.1. Study site and tree species

The experiment was conducted at the Department of Forestry and Environmental Science, Shahjalal University of Science and Technology, Bangladesh, from May 14 to September 16, 2015. Over this period, monthly mean temperatures ranged from 25°C to 32°C, and average monthly rainfall was 295 mm. A 15-m x 3-m raised bed oriented north-south was established and divided into four blocks to enhance drainage and simplify management. We utilized a completely randomized block design, testing eight species (four leguminous and four non-leguminous tree species: Table 1), and three biochar treatments (Control, 10 t/ha, 20 t/ha). This resulted in 96 plants in total, with each block containing a mix of treatments and species (4 replicates x 3 treatments x 8 species).

**Table 1.** Tree species used in this experiment.

| Scientific name | Family | Shade tolerance |
| --- | --- | --- |
| *Acacia auriculiformis* A.Cunn. ex Benth. | Fabaceae (Mimosoideae) | intolerant |
| *Acacia mangium* Willd. | Fabaceae (Mimosoideae) | intolerant |
| *Delonix regia* (Boj. ex Hook.) Raf. | Fabaceae (Caesalpinioideae) | intolerant |
| *Pterocarpus santalinus* L.f. | Fabaceae (Faboideae) | intolerant |
| *Eucalyptus alba* Reinw. ex Blume | Myrtaceae | intolerant |
| *Syzygium cumini* (L.) Skeels. | Myrtaceae | intermediate |
| *Swietenia macrophylla* King. | Meliaceae | intermediate |
| *Melia azedarach* L. | Meliaceae | intolerant |

Seeds of the eight species were sourced from the Bangladesh Forest Department. A hot water treatment was applied to break seed dormancy, and germination occurred in cow-manure-enriched soil within plastic bags. After 45 days, seedlings with similar vigour were transplanted into 3-L pots with a 230 cm² surface area. Biochar-treated pots were filled sequentially with local soil and a 10 cm top layer of biochar-soil mixture (23 g for 10 t/ha and 46 g for 20 t/ha, on a dry mass basis). Incorporation in the upper soil was implemented to simulate surface applications likely in managed forests or restoration applications. No additional fertilization was provided. The biochar dosages used were chosen to be in a range near to somewhat below the optimum of 20-30 t/ha found in dose-response studies (Gale & Thomas, 2019). Biochar was air-dried before mixing with the topsoil layer in each pot. The control group received no biochar, whereas the treatment groups received corresponding biochar quantities. Although it was the monsoon season, during periods of low rainfall an equal amount of deionized/autoclaved water was added to approximate field capacity, a total of two times, to each pot.

### 2.2. Soil

A locally sourced disturbed Dystric Fluvisol sandy loam soil was used in the experiment. Soil samples were collected from each pot following harvest from the uppermost 10 cm, sampled, homogenized, and sieved to <2 mm prior to analysis. The analyses were conducted at the Soil Resource Development Institute (SRDI) in Sylhet, Bangladesh. Soil organic matter (%) was determined by loss-on-ignition using a 6-h combustion time at 600°C in a muffle furnace. Soil pH was determined electrochemically using a 1:2 mixture of soil:deionized water using a pH meter (Kelway MA-78, Kel Instruments, Wycko, NJ, USA). Total N (%) was determined by a semi-micro Kjeldahl method, available P by a Bray-Kurtz I extraction followed by colorimetric analysis, and available K by ammonium acetate extraction followed by flame photometry. Details of soil methods follow Karim et al. (2020). The average (±SE) soil properties were as follows: pH 6.00 ± 0.03, nitrogen content 0.1 ± 0.0%, organic matter content 1.72 ± 0.01%, available phosphorus 59 ± 3 ppm, and potassium 8.8 ± 0.5 ppm, based on three replicates and determined using standard methods.

### 2.3. Biochar

Biochar was produced using a Top-Lit Up-Draft (TLUD) gasifier (Mia et al., 2015) with *Samanea saman* (Jacq.) Merr. wood chips used as the feedstock. Pyrolysis occurred over 15 hours, with a peak temperature of ∼550°C. The resulting biochar had a pH of 8.52 ± 0.02, electrical conductivity (EC) 555 ± 23 dS/m, carbon content 72.07 ± 0.07%, and nitrogen content 1.76 ± 0.014%, averaged from 3 replicate samples (methods follow Karim et al., 2020). Biochar was manually broken into pieces with dimensions < 1 cm (mostly 1-5 mm) prior to soil applications.

### 2.4. Plant measurements

Biweekly measurements of stem height, root collar diameter, leaf count, and leaf length were taken. At harvest, the plants were partitioned into different compartments (main stem, branches, leaves, and roots). Roots were washed with deionized water, nodules were counted (in legume species), and taproot lengths measured. All plant parts were then dried separately at 60°C for at least 48 hours until a constant weight was achieved and weighed to the nearest 0.001 g.

### 2.4. Statistical analysis

Data were analyzed in R (R Core Team, 2024). Initially linear mixed-effects models were implemented using the “lmer” function of the “lme4” package (Bates et al., 2015) and incorporating biochar treatment and species as the main fixed effects, the biochar x species interaction, and random effects of blocks. The random block term was not significant in analyses, so simple two-way ANOVAs with biochar treatment and species as the main effects and the biochar x species interaction term were used. This was followed by one-way ANOVA and Tukey HSD post-hoc tests conducted to compare treatment effects within each species. Agreement with parametric statistical assumptions was assessed using diagnostic plots in conjunction with the Shapiro-Wilk test for normality of residuals and the Levene’s test for homogeneity of variance. Total biomass and that of components (root, leaf, and stem) biomass were right-skewed, so these data were log-transformed prior to analysis. In other cases deviations from assumptions were minor and data transformations were not used. Marginal effects were calculated using the “emmeans” function (Lenth et al., 2021).

To test for treatment effects on height-diameter relationships, we used linear mixed effects models to account for repeated measurements on individual saplings, and assumed linear allometric patterns of the form H = aDb, which are generally found for early growth patterns in tropical trees (Thomas, 1996). Log-transformed height was modeled as a function of log-transformed root collar diameter, species, treatment, and the species x treatment interaction (all fixed effects), and a random individual sapling term. A similar analysis was conducted to determine the relationships between branch number and stem diameter.

### 2.5. Meta-analysis

A meta-analysis of published literature on responses of tropical and sub-tropical trees to biochar was conducted to better assess the generality of findings. Prior meta-analyses of tree responses to biochar (Thomas and Gale, 2015; Juno and Ibáñez, 2021) were used as a starting point. Literature was searched using both ISI Web of Knowledge and Google Scholar, with a cutoff date of June 2024. ISI Web of Knowledge used explicit Boolean operators, while Google Scholar searches used proprietary search algorithms with combinations of 4-5 search terms, and a cutoff of 100 articles scanned per combination. In the case of searches related to post-treatment effects, search terms included descriptions for the set of terms in potential use to describe biochar (including “biochar”, “black carbon”, “char”, “charcoal”, “hydrochar”), in conjunction with (“tropic*” or “sub-tropic*” plus “tree” or “seedling” or “sapling”). Titles and abstracts were scanned for articles that would plausibly present original data on plant growth responses, and tables and figures of those articles were searched for usable data. Articles presenting usable data were themselves read for citations to related articles with potentially usable data. The following additional criteria were used to screen studies: (1) studies examining seed germination or the earliest phases of seedling growth were excluded; (2) species locations or species ranges were located between 25°N and S latitudes. Data from extra-tropical greenhouse studies on tropical and sub-tropical species were also included. The systematic data search located 384 publications, and the overall process (including citation searches) finally yielded a total of 49 publications presenting data useable for meta-analysis.

The response ratio 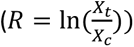 was used as an effect size statistic, where R is the response ratio statistic, X_t_ is the treatment mean and X_c_ is the control mean of the resposnes; pooled R values were inversely weighted by sampling variance. Many studies included cases where multiple biochar treatments (typically different dosages or biochar types) were compared to a single control. Multi-level random effects models were used to account for non-independence within and among studies using the “rma.mv” function in the “metafor” package in R (Viechtbauer, 2010), with a within-study random effect specifying groups of treatments sharing a control, and a between-study random effect accounting for random effects of individual studies (Nakagawa et al., 2023). Models were fitted with restricted maximum likelihood. Data were extracted from tables (or original data) where possible; graphical data were digitized using WebPlotDigitizer (Rohatgi, 2019). Total biomass responses were used as the growth metric where available. Alternative growth measures used included aboveground biomass and the product of tree height and the square of stem diameter, both of which generally scale linearly with total biomass (Kohyama & Hotta, 1990; Deb et al., 2012). In cases where means were presented without error values, standard deviations were imputed from the observed average coefficient of variation observed across studies (Lajeunesse et al., 2013): 𝑆𝐷 = 𝑚𝑒𝑎𝑛 × 𝐶𝑉𝑎𝑣𝑒𝑟𝑎𝑔𝑒. To express response ratios as a percent change, the metric was back-transformed: i.e., percent change = 100 × (exp(R) − 1). Analyses used the metafor package (Viechtbauer, 2010). A similar internal meta-analysis was used to test for differences in legume vs. non-legume growth responses (and biochar dosages) based on data from the present study.

## 3. Results

### 3.1. Soil responses

Biochar additions resulted in variable effects on soil pH, with both notable increases and decreases that depended on the tree species (Fig. 1a). Both the species (F_7,72_ = 13.3; p < 0.001) and species x treatment interaction (F_14,72_ = 6.2; p < 0.001) terms were statistically significant (Table 2). Species showing significant increases in soil pH with biochar additions included *S. macrophylla* and *M. azedarach*; those showing reduced pH in response to biochar additions included *A. mangium*, *D. regia*, and *S. cumini* (Fig. 1a). Although these treatment effects were pronounced, in all cases pH values ranged from ∼6.0 – 6.9 and were thus generally in a near-optimal range.

**Figure 1.**
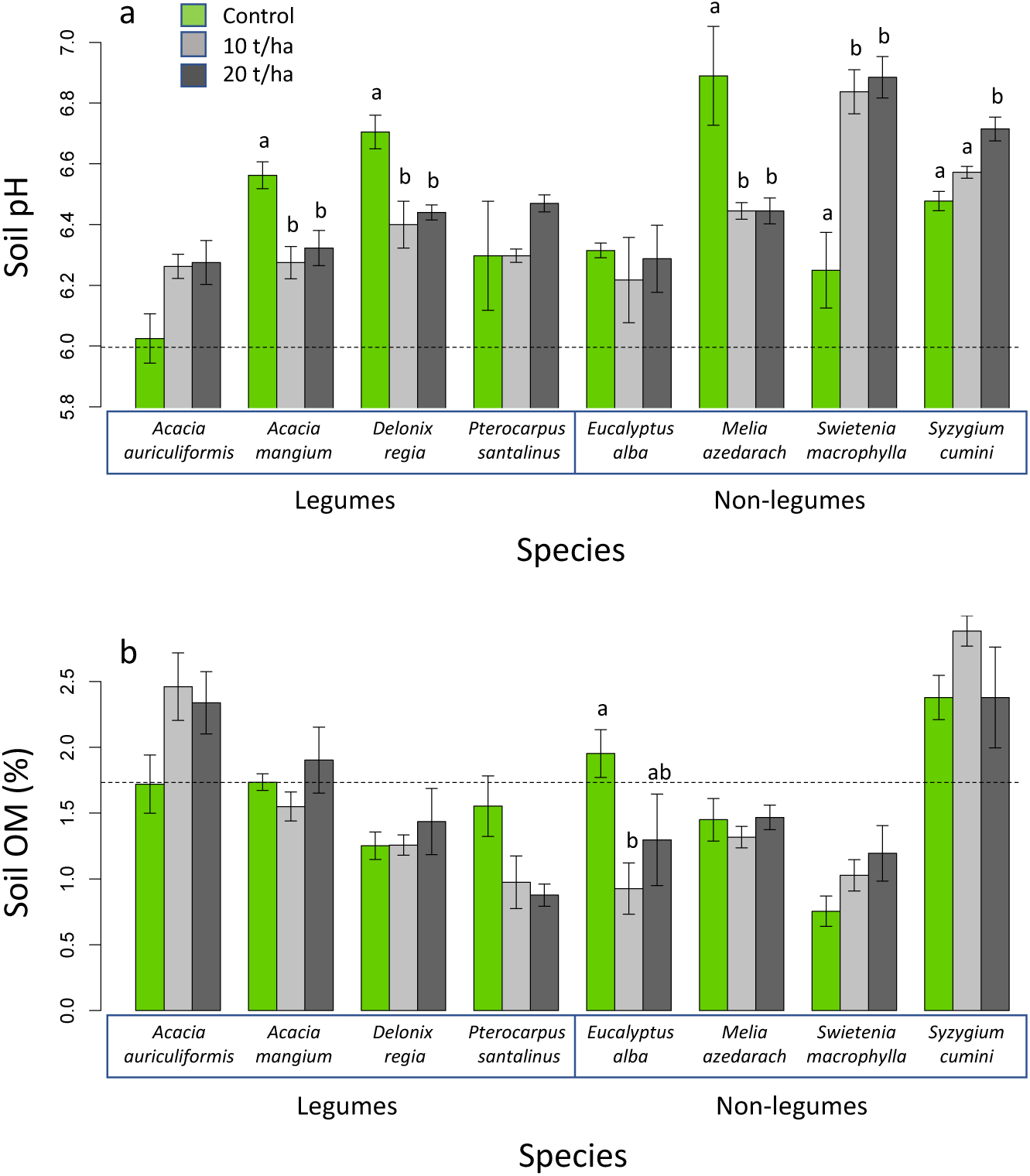
Responses of (a) soil pH, and (b) soil organic matter to biochar addition treatments by species, based on measurements conducted at the end of four-month experimental period. Means are plotted ± 1 SE; dashed lines indicate pre-treatment values. In both cases two-way ANOVA indicates a significant species x biochar treatment interactions (p < 0.001). Separation of means by post-hoc Tukey HSD comparisons (p < 0.05) within each species are indicated by letters.

**Table 2.** ANOVA results for soil parameters measured at the end of four-month experimental period. P-values < 0.05 are listed in bold.

| ANOVA term | d.f. | SS | MS | F | p-value |
| --- | --- | --- | --- | --- | --- |
| pH |  |  |  |  |  |
| Biochar | 2 | 0.07176 | 0.03588 | 1.3915 | 0.2553 |
| Species | 7 | 2.39857 | 0.34265 | 13.2879 | **<0.0001** |
| Biochar x species | 14 | 2.23960 | 0.15997 | 6.2036 | **<0.0001** |
| Residuals | 72 | 1.85665 | 0.02579 |  |  |
| Organic matter |  |  |  |  |  |
| Biochar | 2 | 0.0683 | 0.0341 | 0.2223 | 0.8012 |
| Species | 7 | 23.7921 | 3.3989 | 22.1404 | **<0.0001** |
| Biochar x species | 14 | 5.8741 | 0.4196 | 2.7332 | **0.0028** |
| Residuals | 72 | 11.0530 | 0.1535 |  |  |
| Total N |  |  |  |  |  |
| Biochar | 2 | 0.00014 | 0.000070 | 0.1298 | 0.8784 |
| Species | 7 | 0.07705 | 0.011007 | 20.4784 | **<0.0001** |
| Biochar x species | 14 | 0.01969 | 0.001407 | 2.6171 | **0.0028** |
| Residuals | 72 | 0.03870 | 0.000538 |  |  |
| Available P |  |  |  |  |  |
| Biochar | 2 | 129.37 | 64.684 | 7.0416 | 0.0016 |
| Species | 7 | 327.91 | 46.844 | 5.0995 | **<0.0001** |
| Biochar x species | 14 | 820.46 | 58.605 | 6.3798 | **<0.0001** |
| Residuals | 72 | 661.39 | 9.186 |  |  |
| Available K |  |  |  |  |  |
| Biochar | 2 | 0.057506 | 0.028753 | 15.1084 | **<0.0001** |
| Species | 7 | 0.119399 | 0.017057 | 8.9626 | **<0.0001** |
| Biochar x species | 14 | 0.071360 | 0.005097 | 2.6783 | **0.0033** |
| Residuals | 72 | 0.137025 | 0.001903 |  |  |

Treatment effects on soil OM likewise varied among species, with the ANOVA having significant species (F_7,72_ = 22.1; p < 0.001) and species x treatment interaction (F_14,72_ = 2.7; p = 0.003) terms (Table 2). Although most species showed a trend toward increased soil OM with biochar additions, *E. alba* showed a significant decrease (Fig. 1b).

Biochar effects on macronutrient availability showed pronounced differences among tree species, with a significant species x treatment interaction term for N, P, and K (Table 2). Legume species mostly did not enhance the N status of soils above pre-treatment levels, except for *A. auriculiformis* in biochar-amended soils (Fig. 2a). There were also significant species differences in biochar effects on P and K availability. Biochar additions reduced P availability in *P. santalinus*, *M. azedarach*, and *S. cuminii*, but increased P in *S. macrophylla* (Fig. 2b). Biochar additions also reduced K availability in *D. regina* and *M. azedarach* (Fig. 2c). These species-specific soil nutrient patterns did not have any obvious correspondence with species-specific growth responses to biochar (Fig. 3).

**Figure 2.**
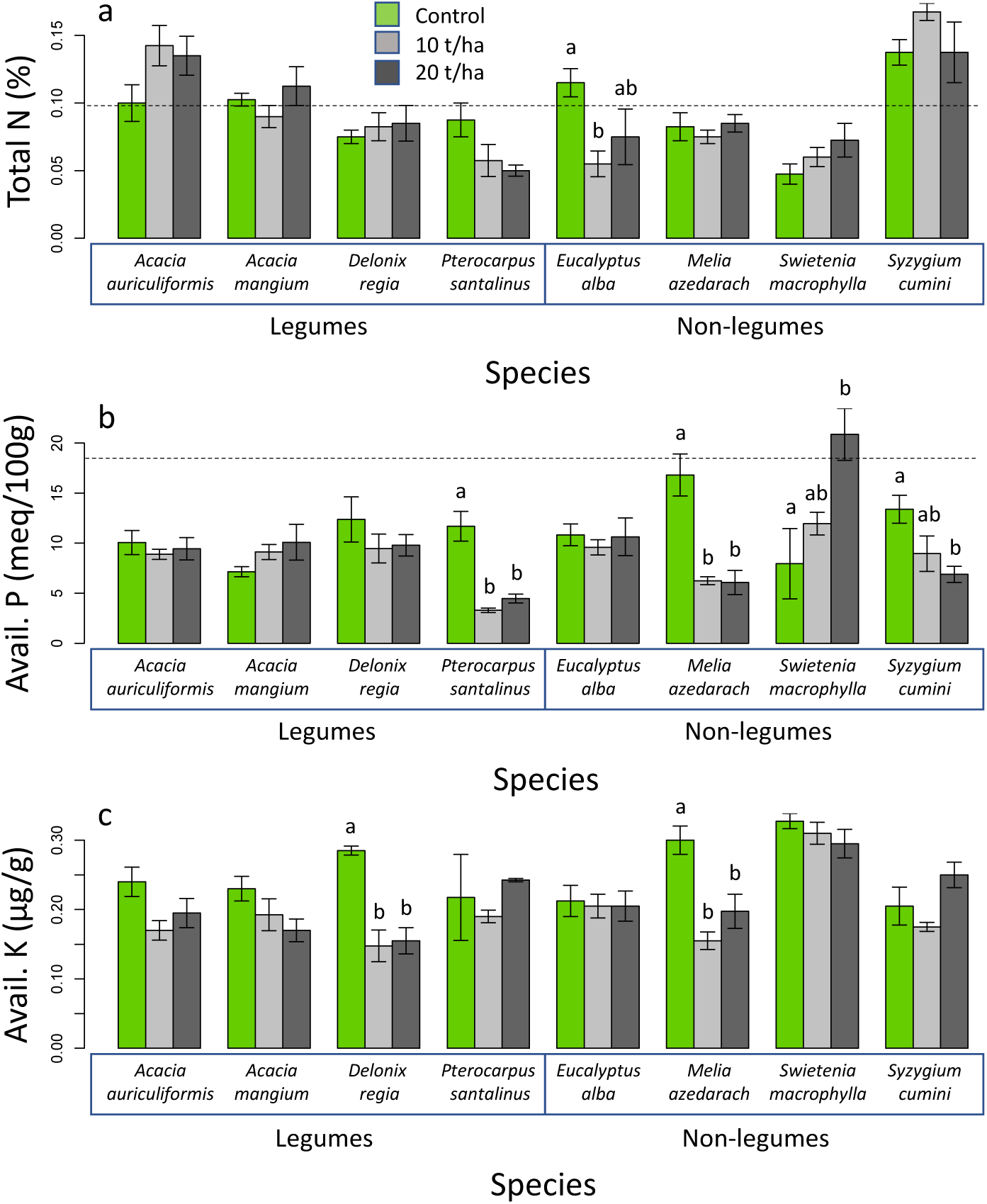
Responses of (a) total soil nitrogen, (b) available phosphorus, and (c) available K, to biochar addition treatments by species, based on measurements at the end of four-month experimental period. Means are plotted ± 1 SE; dashed lines indicate pre-treatment values. (Pre-treatment K values are not shown, as these were considerably higher than post-treatment values). In all three cases two-way ANOVA indicates a significant species x biochar treatment interactions (p < 0.01). Separation of means by post-hoc Tukey HSD comparisons (p < 0.05) within each species are indicated by letters.

**Figure 3.**
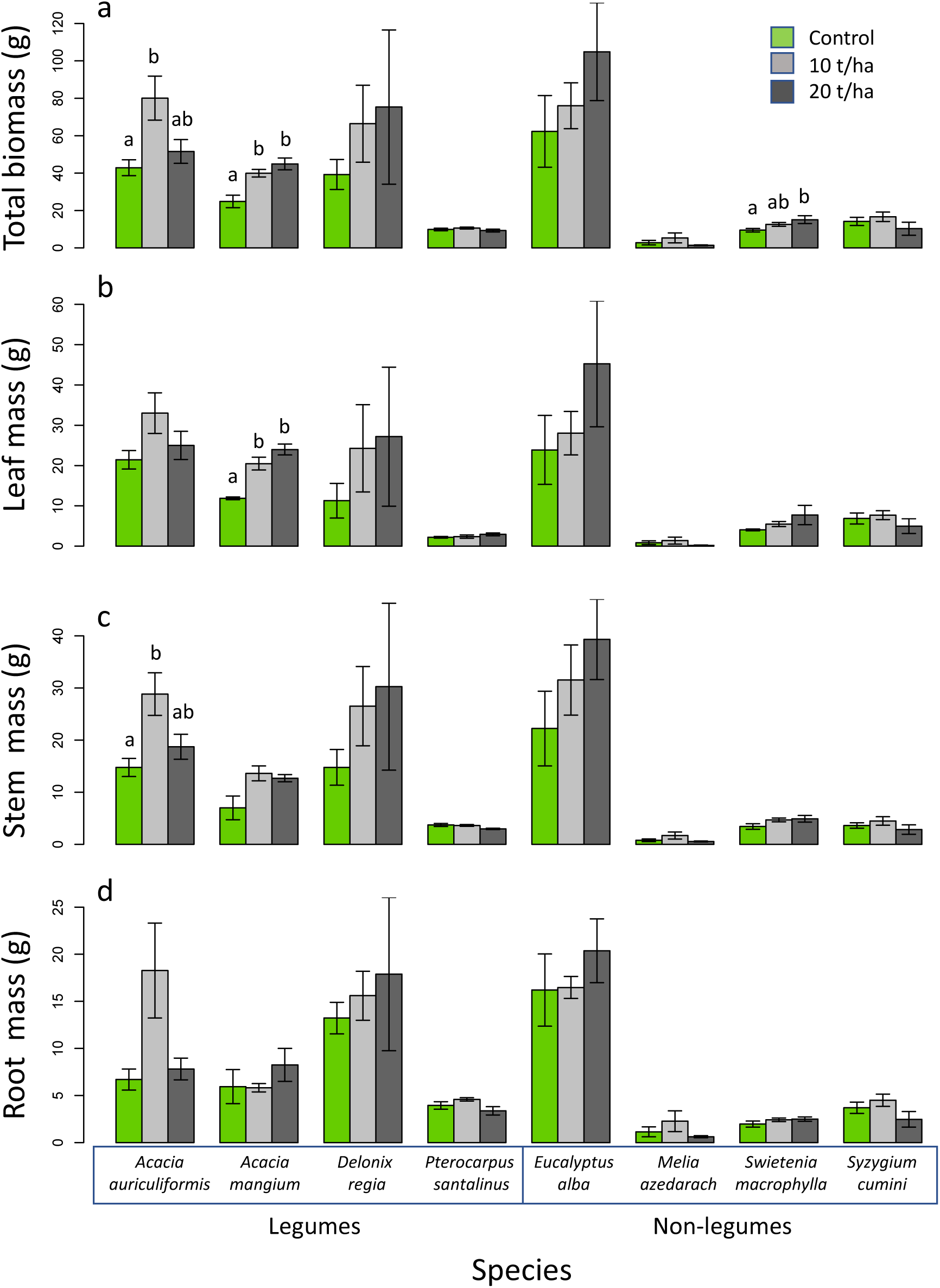
Responses of sapling biomass and biomass components, including (a) total biomass, (b) leaf mass, (c) stem nass, and (d) root mass, to biochar treatments, based on measurements at the end of four-month experimental period. Means are plotted ± 1 SE. Two-way ANOVAs (Table 3) indicate significant biochar treatment effects for total, stem, and root mass (p < 0.05). Separation of means by post-hoc Tukey HSD comparisons (p < 0.05) within each species are indicated by letters.

### 3.2. Sapling growth responses

Biochar additions resulted in strongly enhanced tree growth overall. ANOVA results indicated a significant mean effect term for biochar treatment effects on total biomass (F_2,72_ = 5.1; p = 0.009), stem mass (F_2,72_ = 8.0; p < 0.001), total aboveground biomass (F_2,72_ = 4.9; p = 0.010), and root mass (F_2,72_ = 4.5; p = 0.014), but not leaf mass (F_2,72_ = 1.8; p = 0.174) (Fig. 3; Table 3). Species differences were also significant, but there were no significant treatment x species interactions (Table 3). Tree height and diameter at harvest also showed commensurate positive responses to biochar treatments (Table 3). The overall mean marginal responses were +30% for total biomass (+46% for 10 t/ha; +16% for 20 t/ha), +43% for stem mass (+62% for 10 t/ha; +26% for 20 t/ha), +17% for root mass (+36% for 10 t/ha; +1% for 20 t/ha), and +24% for leaf mass (+47% for 10 t/ha; +5% for 20 t/ha). No mortality was observed.

**Table 3.** ANOVA results for sapling growth parameters measured at the end of four-month experimental period. P-values < 0.05 are listed in bold.

| ANOVA term | d.f. | SS | MS | F | p-value |
| --- | --- | --- | --- | --- | --- |
| Total dry mass |  |  |  |  |  |
| Biochar | 2 | 0.4393 | 0.2196 | 5.0682 | **0.0087** |
| Species | 7 | 21.9754 | 3.1394 | 72.4424 | **<0.0001** |
| Biochar x species | 14 | 0.7628 | 0.0545 | 1.2572 | 0.2555 |
| Residuals | 72 | 3.1202 | 0.0433 |  |  |
| Aboveground dry mass |  |  |  |  |  |
| Biochar | 2 | 0.4922 | 0.2461 | 4.8668 | **0.0104** |
| Species | 7 | 26.178 | 3.7397 | 73.9585 | **<0.0001** |
| Biochar x species | 14 | 0.8799 | 0.0628 | 1.2429 | 0.2649 |
| Residuals | 72 | 3.6407 | 0.0506 |  |  |
| Stem dry mass |  |  |  |  |  |
| Biochar | 2 | 0.7051 | 0.3526 | 7.9915 | **0.0007** |
| Species | 7 | 23.0158 | 3.288 | 74.5299 | **<0.0001** |
| Biochar x species | 14 | 0.8018 | 0.0573 | 1.2982 | 0.2301 |
| Residuals | 72 | 3.1764 | 0.0441 |  |  |
| Leaf dry mass |  |  |  |  |  |
| Biochar | 2 | 0.542 | 0.2709 | 1.7910 | 0.1741 |
| Species | 7 | 37.698 | 5.3855 | 35.6117 | **<0.0001** |
| Biochar x species | 14 | 2.83 | 0.2021 | 1.3367 | 0.2081 |
| Residuals | 72 | 10.888 | 0.1512 |  |  |
| Root dry mass |  |  |  |  |  |
| Biochar | 2 | 0.3771 | 0.1886 | 4.5467 | **0.0138** |
| Species | 7 | 15.0414 | 2.1488 | 51.8153 | **<0.0001** |
| Biochar x species | 14 | 0.7553 | 0.0540 | 1.3009 | 0.2285 |
| Residuals | 72 | 2.9858 | 0.0415 |  |  |
| Stem diameter |  |  |  |  |  |
| Biochar | 2 | 34.84 | 17.418 | 5.0857 | **0.0086** |
| Species | 7 | 1084.33 | 154.904 | 45.2277 | **<0.0001** |
| Biochar x species | 14 | 40.41 | 2.886 | 0.8427 | 0.6217 |
| Residuals | 72 | 246.6 | 3.425 |  |  |
| Height |  |  |  |  |  |
| Biochar | 2 | 4442 | 2220.8 | 8.6844 | **0.0004** |
| Species | 7 | 104979 | 14997 | 58.644 | **<0.0001** |
| Biochar x species | 14 | 4447 | 317.6 | 1.242 | 0.2655 |
| Residuals | 72 | 18413 | 255.7 |  |  |
| Root length |  |  |  |  |  |
| Biochar | 2 | 0.05502 | 0.02751 | 0.9412 | 0.3949 |
| Species | 7 | 1.30568 | 0.18653 | 6.3816 | **<0.0001** |
| Biochar x species | 14 | 0.20663 | 0.01476 | 0.5049 | 0.9229 |
| Residuals | 72 | 2.10448 | 0.02923 |  |  |

Species with notably large responses to biochar additions included both *Acacia* species (with a >80% increase in total biomass for at least one dosage) and *Swietenia macrophylla* (with a 60% increase in the 20 t/ha treatment). Total biomass means by species were larger than controls in all cases at 10 t/ha, and in 6 of 8 cases at 20 t/ha (Fig. 3a). Responses for leaf, stem, and root mass largely mirrored those of total biomass (Fig. 3).

### 3.3. Morphometric and allocation responses

No significant effects of biochar treatments on biomass fractions (i.e., root fraction, leaf fraction, or stem fraction as a proportion of total biomass) were detected on the basis of two-way ANOVAs (Table 4). Root mass per length also did not show a significant treatment effect (Table 4). In all cases there were pronounced differences among species (p < 0.001; Table 4).

**Table 4.**
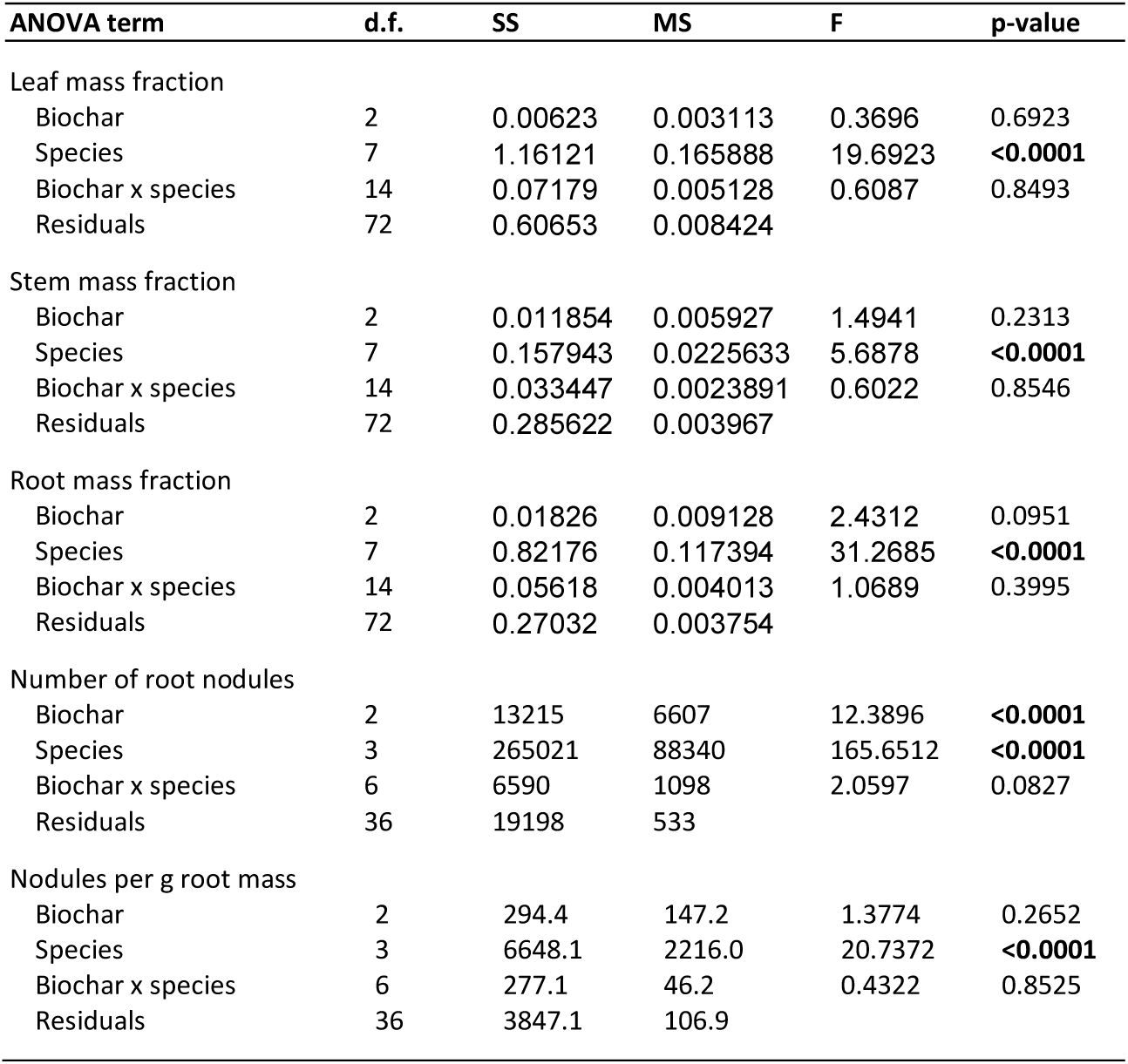
ANOVA results for sapling allocation parameters and root nodulation (in legume species) measured at the end of four-month experimental period. P-values < 0.05 are listed in bold.

Significant biochar treatments effects were found on height-diameter allometric patterns: the linear mixed effects model predicting log(height) had significant terms for log(diameter) (F_1,790_ = 2490.5; p < 0.001), species (F_2,76_ = 110.4; p < 0.001), and biochar treatment (F_7, 76_ = 10.6; p < 0.001), but no significant species x treatment interaction (F_14, 76_ = 1.0; p = 0.498); the random effects term for individual was also significant (p < 0.001). In general, biochar-treated saplings showed increased height values at a given root collar diameter, a pattern most apparent in *D. regia* (p < 0.01) and *S. macrophylla* (p < 0.05) (Fig. 5), with similar trends approaching significance in other species. Based on the fitted model parameters, saplings showed an average increase of 11.1% in height at a given diameter in the 10 t/ha treatment, and a 14.4% increase in the 20 t/ha treatment.

A similar analysis examined potential effects on branch production (i.e., based on log branch number + 1 vs. root collar diameter), but no significant effects were observed (data not shown).

### 3.4. Root nodulation responses in legumes

Biochar additions resulted in increased root nodule production in the four legume species (Table 4; Fig. 5a): the main treatment effect was significant (F_2,36_ = 12.4; p < 0.001), but not the species x treatment interaction (F_6,36_ = 12.4; p < 0.001). Responses were most pronounced in the two *Acacia* species, which also showed the greatest nodule production (Fig. 5). Biochar treatments did not significantly affect the nodulation rate expressed as nodules per g of root tissue (F_2,36_ = 1.4; p = 0.2652), thought the 20 t/ha treatment numerically showed the highest value for three of the four legume species (Fig. 5b). By both metrics nodule production was much greater in the two *Acacia* species than in the other taxa (Fig. 5).

### 3.5. Meta-analyses

Considering only results from the present study, the overall value of the log response ratio R for total biomass is 0.272 ±0.076 (±SE), corresponding to an average 31.3% increase (Z = 3.6; p < 0.001). Values for the two treatment dosages are 0.315 ±0.092 for 10 t/ha (Z = 3.6; p < 0.001) and 0.205 ±0.138 for 20 t/ha (Z = 1.5; p = 0.137). The pooled value for R for total biomass considering legume species is 0.317±0.110 (+37.3%: Z = 2.9; p = 0.004) vs. 0.263 ±0.083 (+30.1%: Z = 3.2; p = 0.002) for non-legumes. Including biochar dosages and legumes vs. non-legumes in the meta-analysis model as moderating factors did not indicate a significant effect of either dosage (Z = 0.724; p = 0.469) or taxonomic status as a legume (Z = 0.640; p = 0.522).

In the broader meta-analysis including prior published studies on tropical and sub-tropical trees, we located a total of 51 publications (Appendix Table A2) presenting useable data that totalled 262 comparisons between control and biochar-amended soils, with 66 comparisons available for legumes (of 23 species in 19 genera) and 173 for non-legume hardwoods (of 37 species in 34 genera including palms), and 21 comparisons for conifers (of 4 species in 3 genera). Averages of reported values for biochar parameters were a peak pyrolysis temperature of 478°C (±78°C SD), pH 8.2 (±1.1), and total C 62% (±17%); these values did not differ significantly among groups (using linear mixed effects models with study included as a random effect). Groups other than conifers show a significant mean increase in biomass in biochar-amended treatments compared to controls: legumes +50.4% (Z = 3.816; p < 0.001), non-legume angiosperms +43.0% (Z = 3.759; p < 0.001), conifers +12.3% (Z = 1.694; p = 0.090), as does the pooled data (+39.3%: Z = 4.600; p < 0.001). Although the mean biomass response is numerically lower in conifers and higher in legumes than the other groups, the inclusion of taxonomic category as a moderator falls short of statistical significance (Q = 1.057; p = 0.589). In all analyses, there was significant residual heterogeneity (Q-tests: p < 0.001). Alternative analyses excluding conifers, excluding data from the present study, excluding data from additions of traditional mound charcoal all yielded similar results (Appendix Table A1).

## 4. Discussion

### 4.1. Patterns in growth responses

Overall, biochar amendments had substantial positive effects on sapling growth in the present study, with a mean biomass increase of 30%. This is generally consistent with meta-analysis results suggesting relatively large growth responses to biochar in tropical trees (Fig. 6; Thomas & Gale, 2015) as is also the case in tropical agricultural systems (Jeffrey et al., 2017), though the responses observed are less than those found in some prior studies (e.g., Sovu et al., 2012; Kayama et al., 2022; Sujeeun & Thomas, 2022). As predicted, the average biomass response observed for legumes was greater than in other species (+37% vs. +30%); however, this difference was not statistically significant. The literature-based meta-analysis gives a much broader representation of species and greater statistical power. This analysis is also consistent with higher biomass responses in legumes (mean +50%) compared to non-legume hardwood species (mean +43%) (Fig. 5), both of which are considerably higher than the mean response for tropical and sub-tropical conifers (mean +12%). However, these statistical comparisons also fall well short of significance due to high variability in responses and relatively low sample sizes in most studies (including the present one).

While the results are suggestive of a generally greater growth response to biochar additions in legumes, there is high variation in responses within groups that obscures any general patterns. Whence this high variability? In some cases very large growth responses reported in the literature are related to exceptional soil conditions. For example, the largest growth responses in the meta-analysis reported here are for native species in the Indian Ocean island of Mauritius in the context of soil impacted by allelopathic strawberry guava (*Psidium cattleyanum*); in this case *Tambourissa peltata* (Monimiaceae) and *Pittosporum senacia* (Pittosporaceae) both showed >40-fold increases in allometric biomass estimates in response to biochar additions (Sujeeun & Thomas, 2022). These exceptional responses are consistent with observations that biochars sorb a wide range of allelochemicals and may also enhance their breakdown, such that biochar additions can completely “rescue” plants suppressed by allelochemicals (Sujeeun & Thomas, 2023). Such a mechanism is obviously not directly related to plant mineral nutrition, and so is not expected to show any systematic relationship with N-fixation. Similarly, large positive effects of biochar on plant performance on contaminated soils can be related to biochar sorption of anthropogenic organic soil contaminants (Ni et al., 2020), potentially toxic elements (Riswan et al., 2016), and sodium salts (Thomas et al., 2013).

Regarding the meta-analysis, its sample size and scope were strongly limited by data reporting practices in the peer-reviewed literature. Many relevant studies did not present means, sample sizes, and/or standard deviations (or data necessary to calculate these) for tree aboveground or total biomass, even in graphical form. This includes numerous examples of recent publications that present “box and whisker” plots that present only medians and inter-quartile plus total data ranges from which it is not possible to calculate means or standard deviations. We were able to compensate for this to some extent by using d2h as a proxy for biomass (Kohyama & Hotta, 1990; Deb et al., 2012) and by using imputed standard deviations (Lajeunesse et al., 2013) for ∼36% of the comparisons. As is a common refrain in the meta-analysis studies, we implore authors to directly publish data or present data in a form that can be used in meta-analyses in future publications. Increased sample sizes within treatments would also aid in detecting patterns: it is notable that the modal sample size found across studies was only N = 3 and median N = 4. There is also a preponderance of pot trials, with only a few multi-year field studies published to date (Table A2).

### 4.2. Mechanism and variable responses within groups

In cases where the mechanism for biochar growth stimulation is mainly related to plant mineral nutrition, the basis for predicting greater growth responses in legumes is their capacity for N-fixation. However, it is important to note that root symbiotic N-fixation exists in tropical woody species outside of the Fabaceae, with rhizobial associations in the angiosperm families Cannabaceae and Zygophyllaceae, cyanobacterial associations in Cycads, and actinorhizal associations in at least some species of the Casuarinaceae, Coriariacea, Elaeagnaceae, Myricaceae, Rhamnaceae, and Rosaceae (Dryadeae tribe) (Tedersoo et al., 2018). We are aware, however, of only one relevant study of a known tropical woody N-fixer that is not a legume: namely, a study of a *Casuarina* species (Mwadalu et al., 2020); in this case, biochar additions alone did not have a positive effect on tree growth. Also, while we confirmed nodulation in the species considered here (Fig. 5), some legumes do not form root nodules (Tedersoo et al., 2018), and the level of root nodulation may not directly correlate with N-fixation (e.g., Quilliam et al., 2013). It is notable that *Acacia mangium* and *A. auriculiformis*, which here showed the largest growth responses to biochar, are both species associated with frequent fire that exhibit putative fire adaptations such as extended seed dormancy and heat-triggered seed germination (Boland et al., 1990; Hegde et al., 2013).

It is generally expected that biochar additions will increase soil pH through a liming effect (Gezahegn et al., 2019) with the potential to increase bioavailability of most nutrients on acidic soils while reducing that of aluminum (Hale et al., 2020, Rios Guayasamín et al., 2024). Here we found that the species effect on soil pH was more pronounced than that of biochar, and also that biochar effects on soil pH varied in both direction and magnitude depending on tree species. There were similar pronounced species x biochar treatment effects on other soil variables examined, including organic matter, and measures of soil N, P, and K (Fig. 1–2; Table 2). Prior studies that have examined responses of multiple species to biochar (e.g., Sovu et al., 2012; Pluchon et al., 2014; Gale et al., 2017; Lefebvre et al., 2019) have not generally considered possible species effects on soil parameters, nor how such effects might be modified by biochar. Mechanisms by which trees can alter soil pH include production of litter or root exudates high in organic acids, stimulation of mineral weathering and oxidation, and ion uptake, in particular of base cations (Alban, 1982; Finzi et al., 1998). In the present short-term experiment, production of root exudates and other forms of rhizodeposition are the most likely mechanism for effects (e.g., Uselman et al. 2000). *Swietenia macrophylla* in combination with biochar stands out as having the largest positive effects on soil pH and available P (Fig. 1). Such interactive effects of biochar and species on soils are an important area for further work in the context of tropical forest restoration and agroforestry. N-fixation may potentially enhance soil N status, and there is some evidence that biochar enhances this effect and can thus enhance overall soil productivity (e.g., Thomas et al., 2019). In the present study, the only species that plausibly showed this pattern was *Acacia auriculiformis* (Fig. 2a).

### 4.3. Effects on tree growth form

In addition to the stimulatory effects of biochar on tree growth, we also found systematic effects on tree growth form and allometry, with an average 11-14% increase in height extension growth at a given stem diameter (Fig. 4). There was no evidence, however, that this response varied among species or was systematically different between legumes and non-legumes. A similar pattern of biochar additions resulting in increased stem elongation relative to diameter has been noted in a few prior studies (Chen et al., 2021; Sifton et al., 2023); including one nursery study of the tropical tree *Bertholletia excelsa* (Damaceno et al., 2019). Freshly produced biochar can release the plant hormone ethylene (Spokas et al., 2010), but ethylene exposure would generally be expected to result in changes in plant morphology similar to thigmomorphogenesis: i.e., reduced stem elongation and increased leaf thickness. Such a response has been noted in a greenhouse study of *Leuceana leucocephala* (Thomas et al., 2019); however, increased stem elongation relative to diameter seems to be the prevailing response to biochar. Increases in height:diameter ratios of plants in response to mineral nutrient fertilization have been widely observed, primarily in relation to N (Thornley, 1999), but also other nutrients such as P (e.g., Brondani et al., 2008). The physiological mechanism by which increased nutrient levels elicit increased allocation to stem elongation is not well resolved, but the response is thought to be functionally related to enhancing the competitive status of plants under high-nutrient conditions (Thornley, 1999). Allometric shifts induced by biochar should receive more research attention and should optimally be incorporated in estimates of biomass and carbon sequestration.

**Figure 4.**
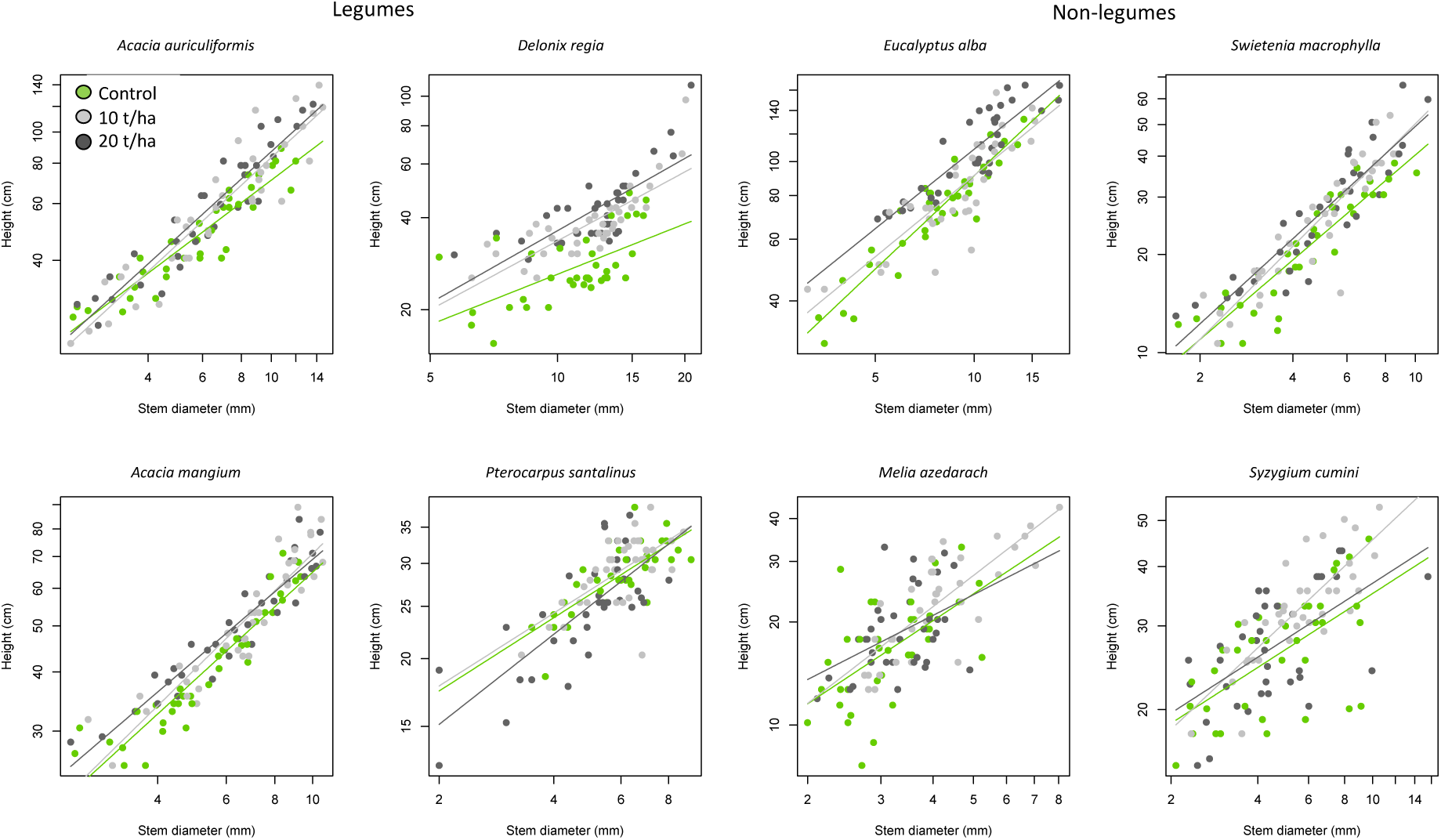
Allometric relationships between sapling height and diameter and their response to biochar treatments. Biochar effects on relationships based on a species-pooled linear mixed model analysis are significant (p < 0.001). Linear regressions for log-transformed variates are shown.

**Figure 5.**
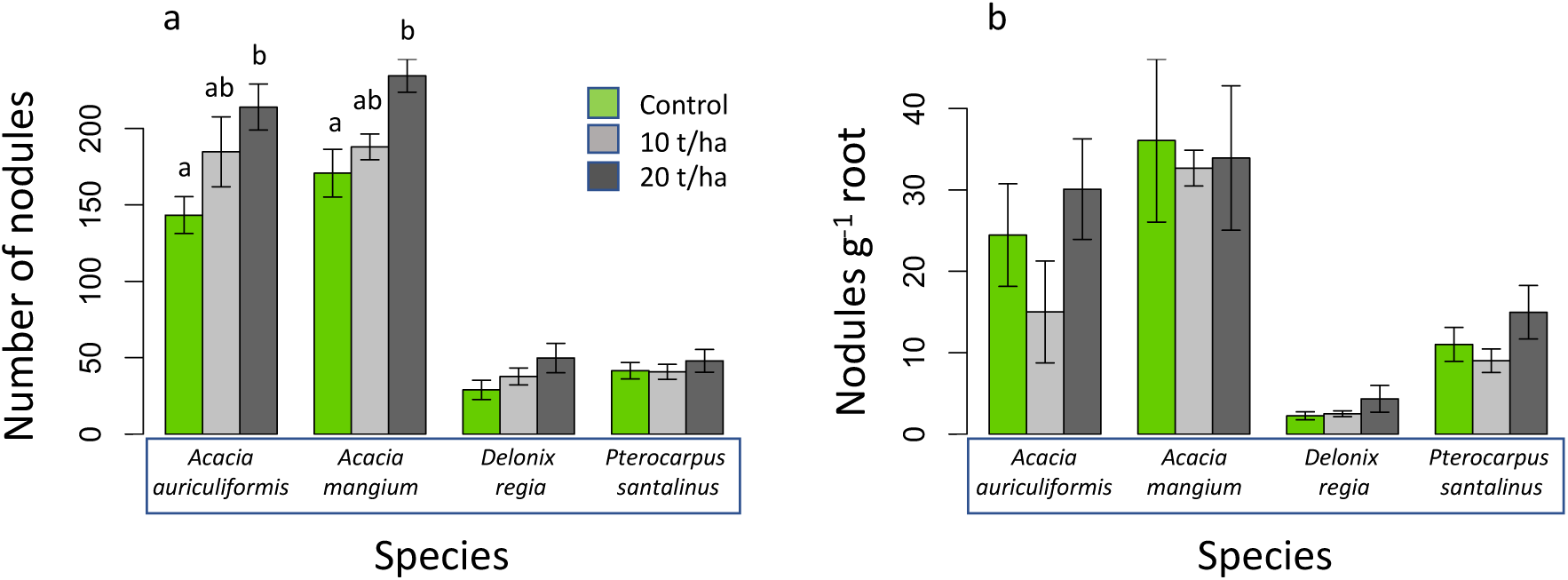
Responses of root nodule production to biochar addtions, expressed as (a) total number of nodules per sapling, (b) nodule number per g root mass. The species-pooled ANOVA indicates a significant biochar main effect for number of nodules. Separation of means by post-hoc Tukey HSD comparisons (p < 0.05) within each species are indicated by letters.

**Figure 6.**
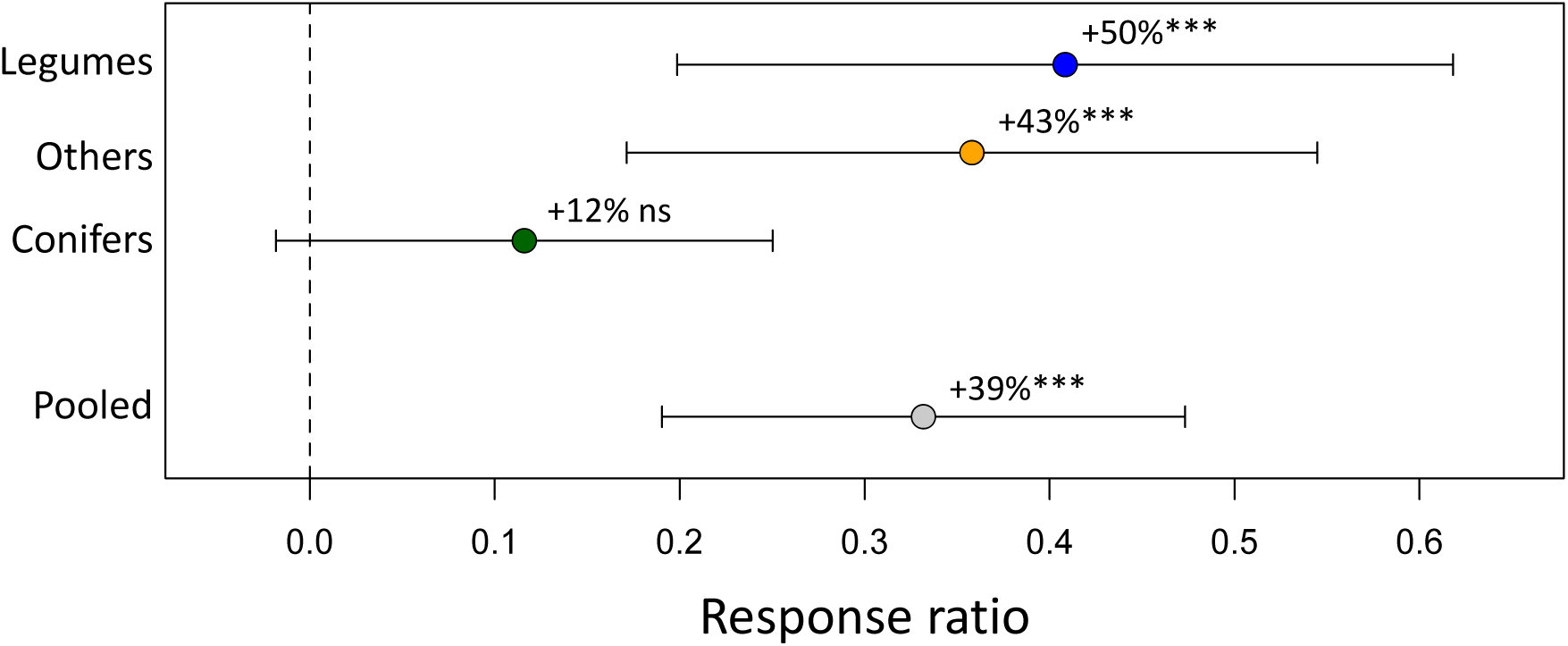
Meta-analysis of biomass responses of tropical and subtropical trees to biochar amendments, based on 38 publications presenting data for 56 tree species. Mean response ratios are plotted ±95% confidence limits, with back-transformed mean responses given with statistical comparisons to zero response: *** p < 0.001.

### 4.4. Conclusion

While this study did not conclusively show that N-fixing leguminous tropical trees respond more strongly to biochar than other species, it reinforces evidence that biochar amendments enhance plant growth and physiological performance in managed ecosystems (Joseph et al., 2021), ranging from conventional agriculture (Schmidt et al., 2021), to forestry (Thomas & Gale, 2015) to urban green infrastructure (Liao et al., 2023). In spite of this vast body of work, one still finds statements in the recent literature to the effect that “biochar seems to have little to no effect on crop yields” (Desjardins et al. 2024, p.12). Of course, it is possible to cite examples of individual studies that find no effect, but the same can be said of virtually any soil amendment or agricultural or silvicultural intervention. The importance of broader survey studies and meta-analyses is highlighted by the considerable variability in responses of tropical trees to biochar, and the multiple mechanisms involved in governing these responses.

It would ultimately be desirable to develop an empirically based decision support tool for use in applied forestry, agroforestry, and forest restoration applications, along the lines of tools being implemented in the context of temperate-zone agriculture (e.g., Latawiec et al., 2017; Phillips et al., 2020). Given the disproportionate importance of tropical forests in global carbon processes and climate feedbacks (Pan et al., 2011; Mitchard, 2018), and the potential for biochar to contribute to global net carbon sinks (Lehmann et al., 2021), development of guidelines for biochar use in tropical forest restoration and plantation forestry should be a global research priority. Well-documented empirical support for the relative benefits of biochar for specific tree species under specific soil conditions is essential in this regard. The present study makes clear that the available empirical data is far from sufficient for tropical trees, in spite of the proliferation of publications in this area. In particular, there is a need for field studies of tree growth responses to biochar on the prevailing low-nutrient status soils of the humid tropics: i.e., Ultisols and Oxisols. There is also an urgent need for additional long-term trials on large trees, as our meta-analysis revealed only four studies of at least 3 years in duration (Table A2). The diversity of tropical trees represents a challenge, but priority should be given to trees of global silvicultural importance (Evans & Turnbull, 2004), such as *Acacia auriculiformis, A. mangium,* and *Swietenia macrophylla*, which stand out in the present study as having exceptionally high growth responses to biochar. More broadly, only ∼1000 tree species make up ∼50% of the stems in tropical forests globally (Cooper et al., 2024). Quantifying biochar responses of a majority of these key species in the global carbon cycle is a realistic objective.

## Appendices

**Figure A1:**
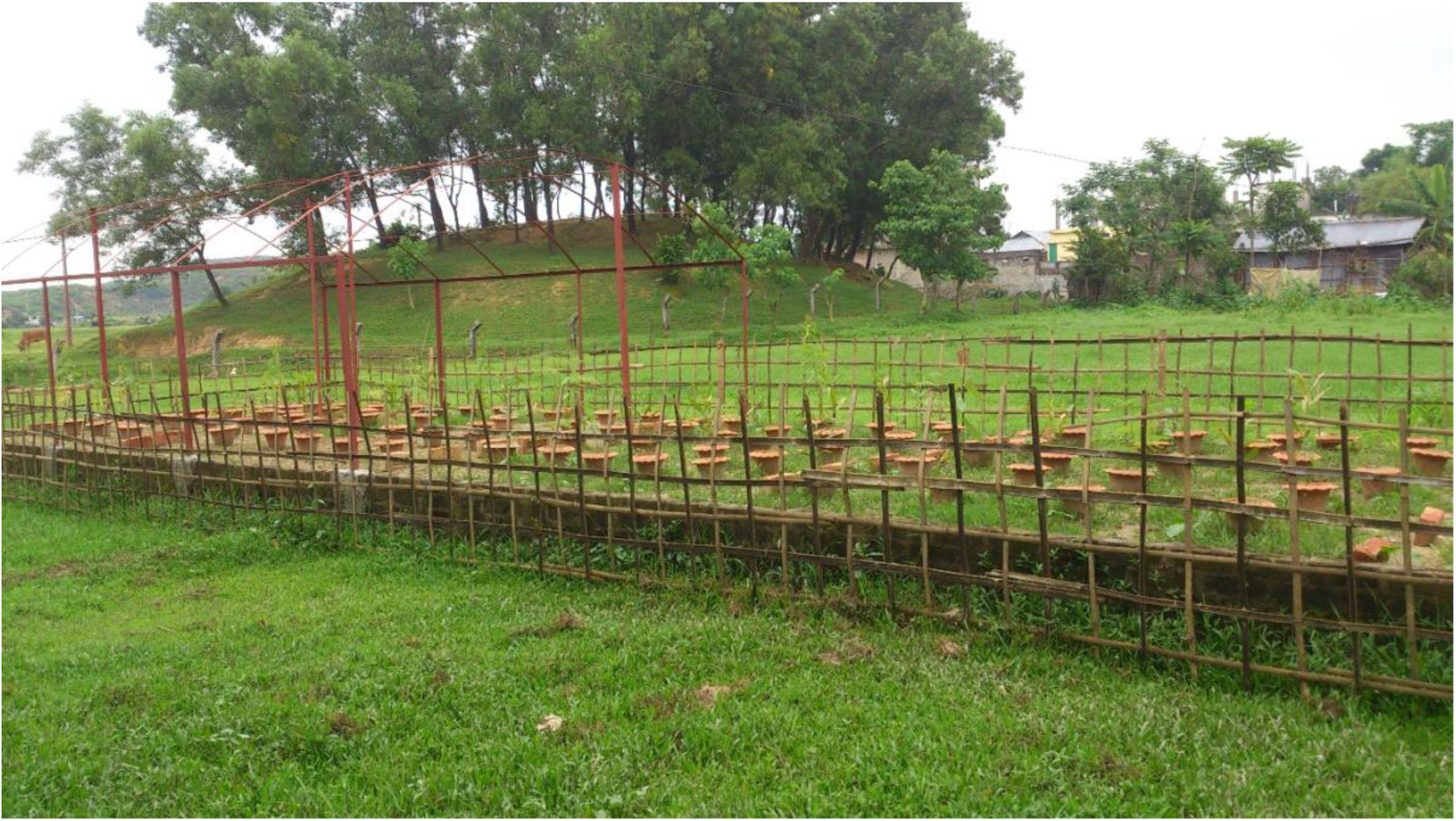
Photo of the experimental study.

**Table A1.** Alternative meta-analysis tests for variation in biochar effects among species groups of tropical and subtropical trees (conifers, legumes, and other hardwoods).

| Data subset | Q | N | p-value |
| --- | --- | --- | --- |
| All data | 1.0574 | 262 | 0.5894 |
| Excluding conifers | 0.0135 | 239 | 0.9076 |
| Excluding current study | 1.2380 | 246 | 0.5385 |
| Excluding traditional mound biochar | 2.0056 | 204 | 0.3669 |
| Excluding studies with imputed errors | 2.6796 | 167 | 0.2619 |

**Table A2.**
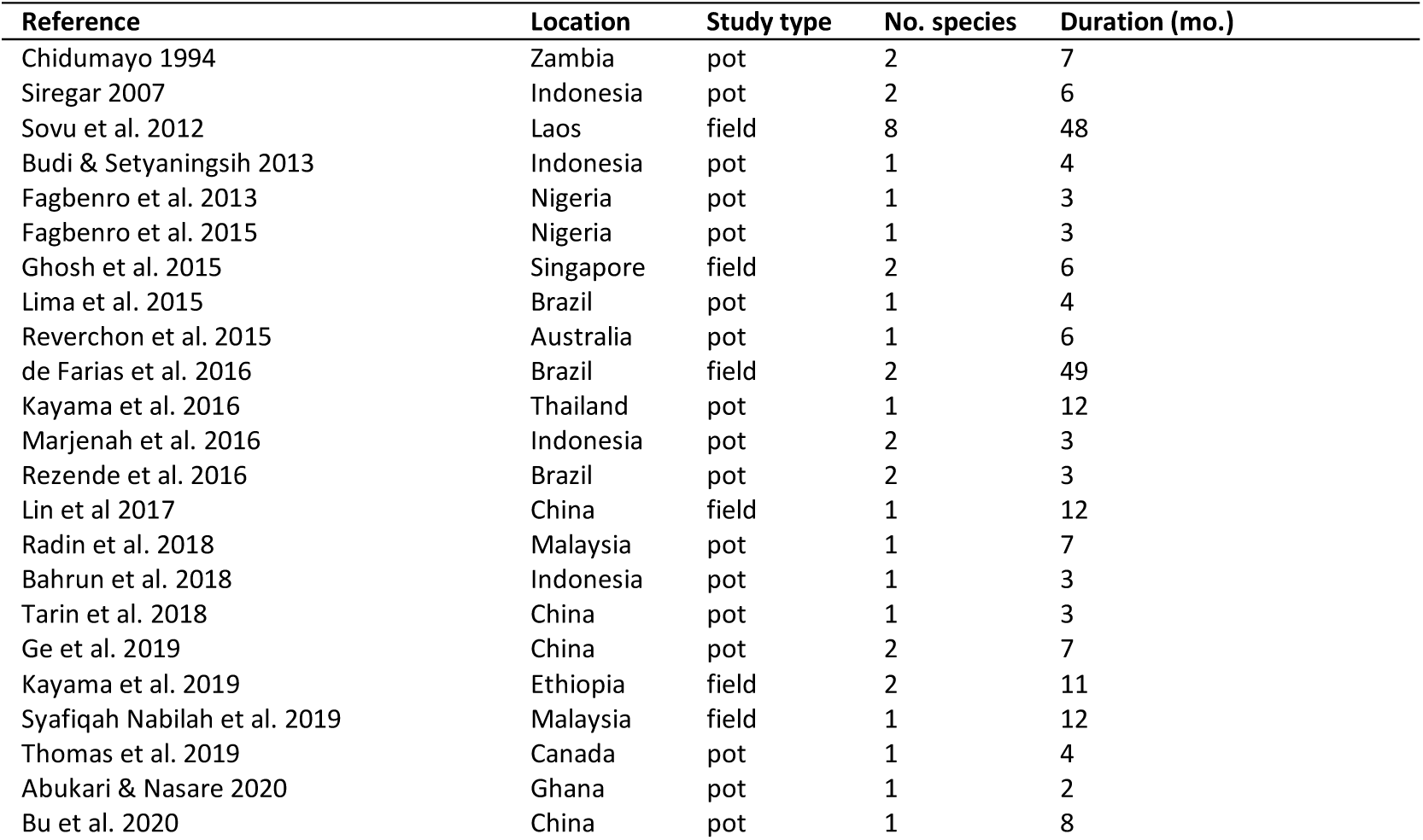
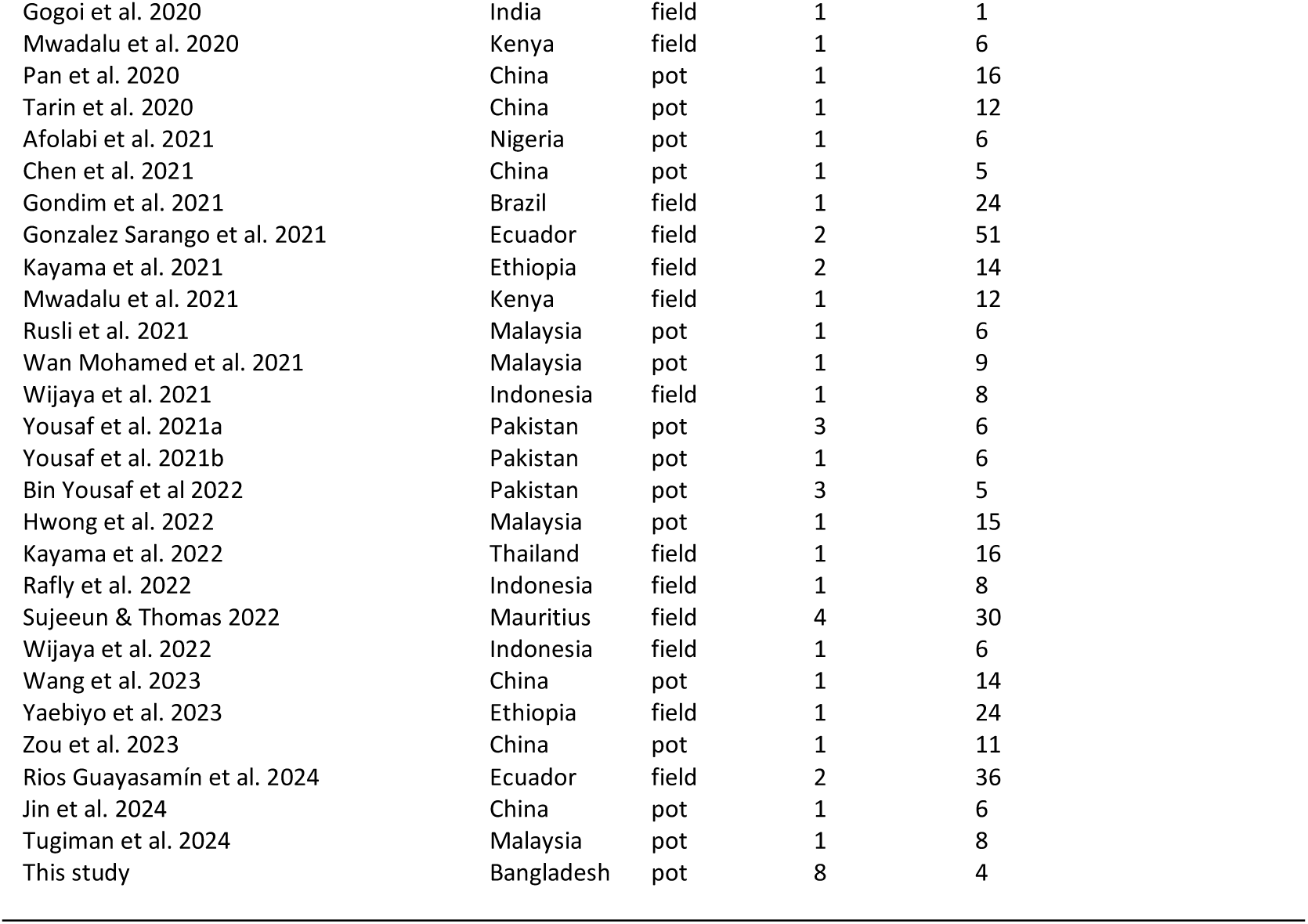
Data sources for literature-based meta-analysis. “Pot” studies include both greenhouse and open-air studies conducted with plants grown in containers.

## Acknowledgements

We acknowledge the logistical support from the Department of Forestry and Environmental Science, Shahjalal University of Science and Technology (SUST), Bangladesh. We are also grateful to Dr. Narayan Saha for his assistance and insights during experiment setup and data collection. We thank Rupon Kumar Nath, Muhammad Ramzan Ali, Sanjoy Bhattacharjee, Halimur Rashid Mannaf, and Tasnima Mukit Riha for help in data collection; nursery assistant Sanatan for support throughout the project; and Rouf Miah of the Bangladesh Forest Department, Sheikhghat, Sylhet, for providing seeds and for guidance with seed germination.

## Data, scripts, code, and supplementary information availability

Orginal data and code used in statistical analyses are available online at https://doi.org/10.5683/SP3/VSRR43.

## References

Abukari A, Nasare NI (2020) Effect of Different Rates of Rice Husk Biochar on the Initial Growth of *Moringa oleifera* under Greenhouse Conditions in the Savannah Ecological Zone of Ghana. Turkish Journal of Agriculture-Food Science and Technology, 8, 13–17. 10.24925/turjaf.v8i1.13-17.2532

Afolabi JO, Abiodun FO, Ojo PA, Ogunwande OA (2021) Influence of watering regimes and bamboo biochar on the growth and biomass partitioning of *Neolamarckia cadamba* (Roxb.) Miq. seedlings on an Alfisol. Ethiopian Journal of Environmental Studies & Management, 14, 515–529. https://ejesm.org/doi/v14i4.10

Alban DH (1982) Effects of nutrient accumulation by aspen, spruce, and pine on soil properties. Soil Science Society of America Journal, 46, 853–861. 10.2136/sssaj1982.03615995004600040037x

Bahrun A, Fahimuddin MY, Rakian TC (2018) Cocoa pod husk biochar reduce watering frequency and increase cocoa seedlings growth. International Journal of Environment, Agriculture and Biotechnology, 3, 1635–1639. 10.22161/ijeab/3.5.9

Bates D, Maechler M, Bolker B, Walker S (2015) Fitting Linear Mixed-Effects Models Using lme4. Journal of Statistical Software, 67, 1–48. 10.18637/jss.v067.i01

Bieser JM, Thomas SC (2019) Biochar and high-carbon wood ash effects on soil and vegetation in a boreal clearcut. Canadian Journal of Forest Research, 49, 1124–1134. 10.1139/cjfr-2019-0039

Bin Yousaf MT, Nawaz MF, Yasin G, Cheng H, Ahmed I, Gul S, Rizwan M, Rehim A, Qi X, Ur Rahman S (2022) Determining the appropriate level of farmyard manure biochar application in saline soils for three selected farm tree species. Plos one, 17, e0265005. 10.1371/journal.pone.0265005

Boland DJ, Pinyopusarerk K, McDonald MW, Jovanovic T, Booth TH (1990) The habitat of *Acacia auriculiformis* and probable factors associated with its distribution. Journal of Tropical Forest Science, 159–180. https://www.jstor.org/stable/43594382

Brondani GE, Silva AJC, Araujo MA., Grossi F, Wendling I, Carpanezzi AA (2008) Phosphorus nutrition in the growth of *Bauhinia forficata* L. seedlings. Acta Scientiarum Agronomy, 30, 665–671. 10.4025/actasciagron.v30i5.787

Bu X, Xue J, Wu Y, Ma W (2020) Effect of biochar on seed germination and seedling growth of *Robinia pseudoacacia* L. in karst calcareous soils. Communications in Soil Science and Plant Analysis, 51, 352–363. 10.1080/00103624.2019.1709484

Budi SW, Setyaningsih L (2013). Arbuscular mycorrhizal fungi and biochar improved early growth of neem (*Melia azedarach* Linn.) seedling under greenhouse conditions. Jurnal Manajemen Hutan Tropika, 19, 103–110. 10.7226/jtfm.19.2.103

Chen H, Chen C, Yu F (2021) Biochar improves root growth of *Sapium sebiferum* (L.) Roxb. container seedlings. Agronomy, 11, 1242. 10.3390/agronomy11061242

Chidumayo EN (1994) Effects of wood carbonization on soil and initial development of seedlings in miombo woodland, Zambia. Forest Ecology and Management, 70, 353–357.

Clough TJ, Condron LM, Kammann C, Müller C (2013) A review of biochar and soil nitrogen dynamics. Agronomy, 3, 275–293. 10.3390/agronomy3020275

Cooper DL, Lewis SL, Sullivan MJ, Prado PI, Ter Steege H, et al. (2024) Consistent patterns of common species across tropical tree communities. Nature, 625, 728–734. 10.1038/s41586-023-06820-z

Damaceno JBD, Lobato ACN, Gama RTD, Oliveira DMD, Falcão NPDS (2019) Biochar as Phosphorus Conditioner in Substrate for Brazil Nut (*Bertholletia excelsa* Humb. & Bonpl.) Seedling Production in the Central Amazon. Journal of Agricultural Science, 11, 383. 10.5539/jas.v11n6p383

Deb JC, Halim MA, Ahmed E (2012) An allometric equation for estimating stem biomass of *Acacia auriculiformis* in the north-eastern region of Bangladesh. Southern Forests: a Journal of Forest Science, 74, 103–113. 10.2989/20702620.2012.701429

de Farias J, Marimon BS, Silva LDCR., Petter FA, Andrade FR, Morandi PS, Marimon-Junior BH (2016) Survival and growth of native *Tachigali vulgaris* and exotic *Eucalyptus urophylla* × *Eucalyptus grandis* trees in degraded soils with biochar amendment in southern Amazonia. Forest Ecology and Management, 368, 173–182.

Desjardins SM, Ter-Mikaelian MT, Chen J (2024) Cradle-to-gate life cycle analysis of slow pyrolysis biochar from forest harvest residues in Ontario, Canada. Biochar, 6, 58. 10.1007/s42773-024-00352-z

Evans J, Turnbull JW (2004) Plantation forestry in the tropics: the role, silviculture, and use of planted forests for industrial, social, environmental, and agroforestry purposes. Oxford University Press.

Fagbenro JA, Oshunsanya SO, Onawumi OA (2013) Effect of Saw Dust Biochar and NPK 15: 15: 15 Inorganic Fertilizer on *Moringa oleifera* Seedlings Grown in an Oxisol. Agrosearch, 13, 57–68. 10.4314/agrosh.v13i1.6

Fagbenro JA, Oshunsanya SO, Oyeleye BA (2015) Effects of gliricidia biochar and inorganic fertilizer on moringa plant grown in an oxisol. Communications in Soil Science and Plant Analysis, 46, 619–626. 10.1080/00103624.2015.1005222

Farhangi-Abriz S, Torabian S, Qin R, Noulas C, Lu Y, Gao S (2021) Biochar effects on yield of cereal and legume crops using meta-analysis. Science of the Total Environment, 775, 145869. 10.1016/j.scitotenv.2021.145869

Farhangi-Abriz S, Ghassemi-Golezani K, Torabian S, Qin R (2022) A meta-analysis to estimate the potential of biochar in improving nitrogen fixation and plant biomass of legumes. Biomass Conversion and Biorefinery, 14, 3293–3303. https://link.springer.com/article/10.1007/s13399-022-02530-0

Finzi AC, Canham CD, Van Breemen N (1998) Canopy tree–soil interactions within temperate forests: species effects on pH and cations. Ecological Applications, 8, 447–454. 10.1890/1051-0761(1998)008[0447:CTSIWT]2.0.CO;2

Fulton W, Gray M, Prahl F, Kleber M (2013) A simple technique to eliminate ethylene emissions from biochar amendment in agriculture. Agronomy and Sustainable Development, 33, 469–474. 10.1007/s13593-012-0118-5

Gale NV, Halim MA, Horsburgh M, Thomas SC (2017) Comparative responses of early-successional plants to charcoal soil amendments. Ecosphere, 8, e01933. 10.1002/ecs2.1933

Gale NV, Sackett TE, Thomas SC (2016) Thermal treatment and leaching biochar alleviates plant growth inhibition from mobile organic compounds. PeerJ, 4, e2385. 10.7717/peerj.2385

Gale NV, Thomas SC (2019) Dose-dependence of growth and ecophysiological responses of plants to biochar. Science of the Total Environment, 658, 1344–1354. 10.1016/j.scitotenv.2018.12.239

Ge X, Yang Z, Zhou B, Cao Y, Xiao W, Wang X, Li MH (2019) Biochar fertilization significantly increases nutrient levels in plants and soil but has no effect on biomass of *Pinus massoniana* (Lamb.) and *Cunninghamia lanceolata* (Lamb.) Hook Saplings during the first growing season. Forests, 10, 612. 10.3390/f10080612

Gei M, Rozendaal DM, Poorter L, Bongers F, Sprent JL, Garner MD, et al. (2018) Legume abundance along successional and rainfall gradients in Neotropical forests. Nature Ecology and Evolution, 2, 1104–1111. 10.1038/s41559-018-0559-6

Gezahegn S, Sain M, Thomas SC (2019) Variation in feedstock wood chemistry strongly influences biochar liming potential. Soil Systems, 3, 26. 10.3390/soilsystems3020026

Ghosh S, Ow LF, Wilson B (2015) Influence of biochar and compost on soil properties and tree growth in a tropical urban environment. International Journal of Environmental Science and Technology, 12, 1303–1310. 10.1007/s13762-014-0508-0

Gogoi L, Gogoi N, Borkotoki B, Kataki R (2020) Efficacy of biochar application on seed germination and early growth of forest tree species in semi-evergreen, moist deciduous forest. Forests, Trees and Livelihoods, 29, 158–175. 10.1080/14728028.2020.1790432

Gondim R, Maia A, Taniguchi C, Muniz C, Araújo TA, de Melo AT, da Silva J (2021) Beneficial effect of biochar on irrigated dwarf-green coconut tree. Atmosphere, 13, 51. 10.3390/atmos13010051

Gonzalez Sarango EM, Valarezo Manosalvas C, Mora M, Villamagua MÁ, Wilcke W (2021) Biochar amendment did not influence the growth of two tree plantations on nutrient-depleted Ultisols in the south Ecuadorian Amazon region. Soil Science Society of America Journal, 85, 862–878. 10.1002/saj2.20227

Hale SE, Nurida NL, Mulder J, Sørmo E, Silvani L, Abiven S, Joseph S, Taherymoosavi S, Cornelissen G (2020) The effect of biochar, lime and ash on maize yield in a long-term field trial in a Ultisol in the humid tropics. Science of the Total Environment, 719, 137455. 10.1016/j.scitotenv.2020.137455

Hegde M, Palanisamy K, Yi JS (2013) *Acacia mangium* Willd. - A fast growing tree for tropical plantation. Journal of Forest and Environmental Science, 29, 1–14. 10.7747/JFS.2013.29.1.1

Heinrich VH, Vancutsem C, Dalagnol R, Rosan TM, Fawcett D, Silva-Junior CH, Cassol HLG, Achard F, Jucker T, Silva CA, House J, Sitch S, Hales TC, Aragão, LEOC (2023) The carbon sink of secondary and degraded humid tropical forests. Nature, 615, 436–442. 10.1038/s41586-022-05679-w

Hwong CN, Sim SF, Kho LK, Teh Y, Harrold LD, Chua KH, Zainal NH (2022) Effects of biochar from oil palm biomass on soil properties and growth performance of oil palm seedlings. Journal of Sustainability Science and Management, 17, 183–200. 10.46754/jssm.2022.4.014

International Biochar Initiative (IBI) (2015) Standardized product definition and product testing guidelines for biochar that is used in soil (IBI Biochar Standards) version 2.1. https://biochar-international.org/characterizationstandard/

Jeffery S, Verheijen FG, van der Velde M, Bastos AC (2011) A quantitative review of the effects of biochar application to soils on crop productivity using meta-analysis. Agriculture, Ecosystems & Environment, 144, 175–187. 10.1016/j.agee.2011.08.015

Jeffery S, Abalos D, Prodana M, Bastos AC, Van Groenigen JW, Hungate BA, Verheijen F (2017) Biochar boosts tropical but not temperate crop yields. Environmental Research Lettters, 12, 053001. 10.1088/1748-9326/aa67bd

Jin X, Liu K, Zhang N, Wu A, Dong L, Wu Q, Zhao M, Li Y, Wang Y (2024). The combined application of arbuscular mycorrhizal fungi and biochar improves the Cd tolerance of *Cinnamomum camphora* seedlings. Rhizosphere, 100939. 10.1016/j.rhisph.2024.100939

Joseph S, Cowie AL, Van Zwieten L, Bolan N, Budai A, Buss W, Cayuela ML, Graber ER, Ippolito JA, Kuzyakov Y, Luo Y, Ok YS, Palansooriya KN, Shepherd J, Stephens S, Weng Z, Lehmann J (2021) How biochar works, and when it doesn’t: A review of mechanisms controlling soil and plant responses to biochar. GCB Bioenergy, 13, 1731–1764. 10.1111/gcbb.12885

Juno E, Ibáñez I (2021) Biochar application and soil transfer in tree restoration: A meta-analysis and field experiment. Ecological Restoration, 39, 158–167. 10.3368/er.39.3.158

Karim MR, Halim MA, Gale NV, Thomas SC (2020) Biochar effects on soil physiochemical properties in degraded managed ecosystems in northeastern Bangladesh. Soil Systems, 4, 69. 10.3390/soilsystems4040069

Kayama M, Abebe B, Birhane E (2021) Effects of Biochar on the Growth of *Vachellia etbaica* and *Faidherbia albida* Planted in Tigray, Northern Ethiopia. Japan Agricultural Research Quarterly: JARQ, 55, 367–378. 10.6090/jarq.55.367

Kayama M, Nimpila S, Hongthong S, Yoneda R, Wichiennopparat W, Himmapan W, et al. (2016) Effects of bentonite, charcoal and corncob for soil improvement and growth characteristics of teak seedling planted on acrisols in northeast Thailand. Forests, 7, 36. 10.3390/f7020036

Kayama M, Nimpila S, Hongthong S, Yoenda R, Himmapan, W, Noda I (2022) Effects of biochar on the early growth characteristics of teak seedlings planted in sandy soil in northeast Thailand. Bulletin of the Forestry and Forest Products Research Institute, 21, 73–81. 10.20756/ffpri.21.1_73

Kayama M, Takenaka K, Abebe B, Birhane E (2019) Effects of biochar on the growth of *Olea europaea* subsp. *cuspidata* and *Dodonaea angustifolia* planted in Tigray, northern Ethiopia. Journal of the Japanese Society of Revegetation Technology, 45, 115–120. 10.7211/jjsrt.45.115

Kochanek J, Long RL, Lisle AT, Flematti GR (2016) Karrikins identified in biochars indicate post-fire chemical cues can influence community diversity and plant development. PloSOne, 11, e0161234. 10.1371/journal.pone.0161234

Kohyama T, Hotta M (1990) Significance of allometry in tropical saplings. Functional Ecology, 4, 515–521. 10.2307/2389319

Lajeunesse MJ, Koricheva J, Gurevitch J, Mengersen K (2013) Recovering missing or partial data from studies: a survey of conversions and imputations for meta-analysis. In: Koricheva J, Gurevitch J, Mengersen K (eds) Handbook of meta-analysis in ecology and evolution. Princeton University Press, Princeton, pp. 195–206. 10.1515/9781400846184-015

Latawiec AE, Peake L, Baxter H, Cornelissen G, Grotkiewicz K, Hale S, Królczyk JB, Kubon M, Łopatka A, Medynska-Juraszek A, Reid BJ, Siebielec G, Sohi SP, Spiak Z, Strassburg BB (2017) A reconnaissance-scale GIS-based multicriteria decision analysis to support sustainable biochar use: Poland as a case study. Journal of Environmental Engineering and Landscape Management, 25, 208–222. 10.3846/16486897.2017.1326924

Lefebvre D, Román-Dañobeytia F, Soete J, Cabanillas F, Corvera R, Ascorra C, Fernandez LE, Silman M (2019) Biochar effects on two tropical tree species and its potential as a tool for reforestation. Forests, 10, 678. 10.3390/f10080678

Lehmann J, Joseph S (2015) Biochar for environmental management: an introduction. In: Lehmann J, Joseph S (eds) Biochar for Environmental Management: Science, Technology and Implementation, 2nd Edn. Earthscan, London, pp. 1–14.

Lehmann J, Cowie A, Masiello CA, Kammann C, Woolf D, Amonette JE, Cayuela ML, Camps-Arbestain M, Whitman T (2021) Biochar in climate change mitigation. Nature Geoscience, 14, 883–892. 10.1038/s41561-021-00852-8

Lenth R, Singmann H, Love J, Buerkner P, Herve M (2021) Emmeans: estimated marginal means, aka least-squares means (R package version 1.5. 1.). The Comprehensive R Archive Network.

Liao W, Halim MA, Kayes I, Drake JA, Thomas SC (2023) Biochar benefits green infrastructure: global meta-analysis and synthesis. Environmental Science & Technology, 57, 15475–15486. 10.1021/acs.est.3c04185

Lima SL, Tamiozzo S, Palomino EC, Petter FA, Marimon-Junior BH (2015) Interactions of biochar and organic compound for seedlings production of *Magonia pubescens* A. St.-Hil. Revista Árvore, 39, 655–661. 10.1590/0100-67622015000400007

Lin Z, Liu Q, Liu G, Cowie AL, Bei Q, Liu B, et al. (2017) Effects of different biochars on *Pinus elliottii* growth, N use efficiency, soil N_2_O and CH_4_ emissions and C storage in a subtropical area of China. Pedosphere, 27, 248–261. 10.1016/S1002-0160(17)60314-X

Liu L, Wang Y, Yan X, Li J, Jiao N, Hu S (2017) Biochar amendments increase the yield advantage of legume-based intercropping systems over monoculture. Agriculture Ecosystems and Environment, 237, 16–23. 10.1016/j.agee.2016.12.026

Liu X, Zhang A, Ji C, Joseph S, Bian R, Li L, Pan G (2013) Biochar’s effect on crop productivity and the dependence on experimental conditions—a meta-analysis of literature data. Plant and Soil, 373, 583–594. 10.1007/s11104-013-1806-x

Marjenah M, Kiswanto K, Purwanti S, Sofyan FPM (2016) The effect of biochar, cocopeat and sawdust compost on the growth of two dipterocarps seedlings. Nusantara Bioscience, 8, 39–44. 10.13057/nusbiosci/n080108

Mia S, Dijkstra FA, Singh B (2018) Enhanced biological nitrogen fixation and competitive advantage of legumes in mixed pastures diminish with biochar aging. Plant and Soil, 424, 639–651. 10.1007/s11104-018-3562-4

Mia S, Uddin N, Hossain SAM, Amin R, Mete FZ, Hiemstra T (2015) Production of Biochar for Soil Application: A Comparative Study of Three Kiln Models. Pedosphere, 25, 696–702. 10.1016/S1002-0160(15)30050-3

Mitchard ET (2018) The tropical forest carbon cycle and climate change. Nature, 559, 527–534. 10.1038/s41586-018-0300-2

Mwadalu RU, Mochoge B, Danga B (2020) Effects of biochar and manure on soil properties and growth of *Casuarina equisetifolia* seedlings at the coastal region of Kenya. Scientific Research and Essays, 15, 52–63. 10.5897/SRE2020.6684

Mwadalu R, Mochoge B, Danga B (2021) Assessing the potential of biochar for improving soil physical properties and tree growth. International Journal of Agronomy, 2021, 1–12. 10.1155/2021/6000184

Nakagawa S, Yang Y, Macartney EL, Spake R, Lagisz M (2023) Quantitative evidence synthesis: a practical guide on meta-analysis, meta-regression, and publication bias tests for environmental sciences. Environmental Evidence, 12, 8. 10.1186/s13750-023-00301-6

Nguyen TTN, Xu CY, Tahmasbian I, Che R, Xu Z, Zhou X, Wallace HM, Bai SH (2017) Effects of biochar on soil available inorganic nitrogen: a review and meta-analysis. Geoderma, 288, 79–96. 10.1016/j.geoderma.2016.11.004

Ni N, Kong D, Wu W, He J, Shan Z, Li J, Dou Y, Zhang Y, Song Y, Jiang X (2020) The Role of Biochar in Reducing the Bioavailability and Migration of Persistent Organic Pollutants in Soil–Plant Systems: A Review. Bulletin of Environmental Contamination and Toxicology, 104, 157–165. 10.1007/s00128-019-02779-8

Ogunwole AA, Ogunwole OD (2023) Effects of Water Application Rates and Sawdust Biochar on the Physicochemical Properties of Soil and Performance of Five Tree Species Used in Urban Landscaping in Ondo, Nigeria. Cities and the Environment (CATE), 16, 7. 10.15365/cate.2023.160207

Pan L, Xu F, Mo H, Corlett RT, Sha L (2021) The potential for biochar application in rubber plantations in Xishuangbanna, Southwest China: a pot trial. Biochar, 3, 65–76. 10.1007/s42773-020-00072-0

Pan Y, Birdsey RA, Fang J, Houghton R, Kauppi PE, Kurz WA, Phillips OL, Shvidenko A, Lewis SL, Canadell JG, Ciais P, Jackson RB, Pacala SW, McGuire AD, Piao S, Rautiainen A, Sitch S, Hayes D (2011) A large and persistent carbon sink in the world’s forests. Science, 333, 988–993. 10.5194/nhess-24-445-2024

Phillips CL, Light SE, Lindsley A, Wanzek TA, Meyer KM, Trippe KM (2020) Preliminary evaluation of a decision support tool for biochar amendment. Biochar 2, 93–105. 10.1007/s42773-020-00037-3

Pluchon N, Gundale MJ, Nilsson MC, Kardol P, Wardle DA (2014) Stimulation of boreal tree seedling growth by wood-derived charcoal: effects of charcoal properties, seedling species and soil fertility. Functional Ecology, 28, 766–775. 10.1111/1365-2435.12221

Powers JS, Marín-Spiotta E (2017) Ecosystem processes and biogeochemical cycles in secondary tropical forest succession. Annual Review of Ecology Evolution and Systematics, 48, 497–519. 10.1146/annurev-ecolsys-110316-022944

Quilliam RS, DeLuca TH, Jones DL (2013) Biochar application reduces nodulation but increases nitrogenase activity in clover. Plant and Soil, 366, 83–92. 10.1007/s11104-012-1411-4

R Core Team (2024) R: A Language and Environment for Statistical Computing. R Foundation for Statistical Computing, Vienna, Austria.

Radin R, Abu Bakar R, Ishak CF, Ahmad SH, Tsong LC (2018) Biochar-compost mixture as amendment for improvement of polybag-growing media and oil palm seedlings at main nursery stage. International Journal of Recycling of Organic Waste in Agriculture, 7, 11–23. 10.1007/s40093-017-0185-3

Rafly NM, Riniarti M, Hidayat W, Prasetia H, Wijaya BA, Niswati A, Hasanudin U, Banuwa IS (2022) The Effect of the Application of Biochar from Oil Palm Empty Bunches on the Growth of *Falcataria moluccana*. Journal of Tropical Upland Resources, 4, 1–10. 10.23960/jtur.vol4no1.2022.124

Reverchon F, Yang H, Ho TY, Yan G, Wang J, Xu Z, et al. (2015) A preliminary assessment of the potential of using an acacia—biochar system for spent mine site rehabilitation. Environmental Science and Pollution Research, 22, 2138–2144. 10.1007/s11356-014-3451-1

Rezende FA, Santos VAHFD, Maia CMBDF, Morales MM (2016) Biochar in substrate composition for production of teak seedlings. Pesquisa Agropecuária Brasileira, 51, 1449–1456. 10.1590/S0100-204X2016000900043

Riniarti M, Hidayat W, Prasetia H, Niswati A, Hasanudin U, Banuwa, Yoo J, Kim S, Lee S (2021) Using two dosages of biochar from shorea to improve the growth of *Paraserianthes falcataria* seedlings. IOP Conference Series: Earth and Environmental Science, 749, 012049. 10.1088/1755-1315/749/1/012049

Rios Guayasamín PD, Smith SM, Thomas SC (2024) Biochar effects on NTFP-enriched secondary forest growth and soil properties in Amazonian Ecuador. Journal of Environmental Management, 350, 119068. 10.1016/j.jenvman.2023.119068

Rizwan M, Ali S, Qayyum MF, Ibrahim M, Zia-ur-Rehman M, Abbas T, Ok YS (2016) Mechanisms of biochar-mediated alleviation of toxicity of trace elements in plants: a critical review. Environmental Science and Pollution Research, 23, 2230–2248. 10.1007/s11356-015-5697-7

Robertson SJ, Rutherford PM, Lopez-Gutierrez JC, Massicotte HB (2012). Biochar enhances seedling growth and alters root symbioses and properties of sub-boreal forest soils. Canadian Journal of Soil Science, 92, 329–340. 10.4141/cjss2011-066

Rohatgi A (2019) WebPlotDigitizer (Version 3) [Software]. Retrieved from https://apps.automeris.io/wpd4/

Rondon MA, Lehmann J, Ramírez J, Hurtado M (2007) Biological nitrogen fixation by common beans (*Phaseolus vulgaris* L.) increases with bio-char additions. Biology and Fertility of Soils, 43, 699–708. 10.1007/s00374-006-0152-z

Rusli LS, Osman N, Abdullah R, Yaacob JS, Seow AH (2021) Effects of palm kernel biochar on the physiological responses and root profiles of sendudok (*Melastoma malabathricum* L.) grown on acidic soil. Applied Ecology and Environmental Research, 19, 2887–2903. 10.15666/aeer/1904_28872903

Schmidt HP, Kammann C, Hagemann N, Leifeld J, Bucheli TD, Sánchez Monedero MA, Cayuela ML (2021) Biochar in agriculture–A systematic review of 26 global meta-analyses. GCB Bioenergy, 13, 1708–1730. 10.1111/gcbb.12889

Sifton MA, Lim P, Smith SM, Thomas SC (2022) Interactive effects of biochar and N-fixing companion plants on growth and physiology of *Acer saccharinum*. Urban Forestry and Urban Greening, 74, 127652. 10.1016/j.ufug.2022.127652

Sifton MA, Smith SM, Thomas SC (2023). Biochar-biofertilizer combinations enhance growth and nutrient uptake in silver maple grown in an urban soil. PloSOne, 18, e0288291. 10.1371/journal.pone.0288291

Siregar CA (2007) Effect of charcoal application on the early growth stage of *Acacia mangium* and *Michelia montana*. Indonesian Journal of Forestry Research, 4, 19–30.

Spokas KA, Baker JM, Reicosky DC (2010) Ethylene: potential key for biochar amendment impacts. Plant and Soil, 333, 443–452. 10.1007/s11104-010-0359-5

Spokas KA, Cantrell KB, Novak JM, Archer DW, Ippolito JA, Collins HP, Boateng AA, Lima IM, Lamb MC, McAloon AJ, Lentz RD, Nichols KA (2012) Biochar: a synthesis of its agronomic impact beyond carbon sequestration. Journal of Environmental Quality, 41, 973–989. 10.2134/jeq2011.0069

Sovu MT, Savadogo P, Odén PC (2012) Facilitation of forest landscape restoration on abandoned swidden fallows in Laos using mixed-species planting and biochar application. Silva Fennica, 46, 39–51. 10.14214/sf.444

Sujeeun L, Thomas SC (2022) Biochar rescues native trees in the biodiversity hotspot of Mauritius. Forests, 13, 277. 10.3390/f13020277

Sujeeun L, Thomas SC (2023) Biochar: A Tool for Combatting Both Invasive Species and Climate Change. Pages 367-393 in: Tripathi S, Bhadouria R, Srivastava P, Singh R, Batish DR (eds) Plant Invasions and Global Climate Change. Springer, Singapore. 10.1007/978-981-99-5910-5_16

Syafiqah Nabilah SB, Fazwa F, Norhayati S, Jeyanny V, Mohd Zaki, Mohd Asri L, Samsuri TH (2019) Effect of NPK Fertilizer, Biochar and Compost on the Growth of Citrus hystrix. Transactions of the Malaysian Society of Plant Physiology, 26, 47–52.

Tamme R, Pärtel M, Kõljalg U, Laanisto L, Liira J, Mander Ü, Moora M, Niinemets Ü, Öpik M, Ostonen I, Tedersoo L, Zobel M (2021) Global macroecology of nitrogen-fixing plants. Global Ecology and Biogeography, 30, 514–526. 10.1111/geb.13236

Tarin MWK, Fan L, Tayyab M, Sarfraz R, Chen L, He T, Rong J, Chen L, Zheng Y (2018) Effects of bamboo biochar amendment on the growth and physiological characteristics of *Fokienia hodginsii*. Applied Ecology & Environmental Research, 16, 8055–8074. 10.15666/aeer/1606_80558074

Tarin MWK, Fan L, Shen L, Lai J, Li J, Deng Z, Chen L, He T, Rong J, Zheng, Y. (2020) Rice straw biochar impact on physiological and biochemical attributes of *Fokienia hodginsii* in acidic soil. Scandinavian Journal of Forest Research, 35, 59–68. 10.1080/02827581.2020.1731591

Tedersoo L, Laanisto L, Rahimlou S, Toussaint A, Hallikma T, Pärtel M (2018). Global database of plants with root-symbiotic nitrogen fixation: NodDB. Journal of Vegetation Science, 29, 560–568. 10.1111/jvs.12627

Thomas SC (1996) Asymptotic height as a predictor of growth and allometric characteristics in Malaysian rain forest trees. American Journal of Botany, 83, 556–566. 10.1002/j.1537-2197.1996.tb12739.x

Thomas SC, Frye S, Gale N, Garmon M, Launchbury R, Machado N, Melamed S, Murray J, Petroff A Winsborough C (2013) Biochar mitigates negative effects of salt additions on two herbaceous plant species. Journal of Environmental Management, 129, 62–68. 10.1016/j.jenvman.2013.05.057

Thomas SC, Gale N (2015) Biochar and forest restoration: a review and meta-analysis of tree growth responses. New Forests, 46, 931–946. 10.1007/s11056-015-9491-7

Thomas SC, Halim MA, Gale NV, Sujeeun L (2019) Biochar enhancement of facilitation effects in agroforestry: early growth and physiological responses in a maize-leucaena model system. Agroforestry Systems, 93, 2213–2225. 10.1007/s10457-018-0336-1

Thornley JH (1999) Modelling stem height and diameter growth in plants. Annals of Botany, 84, 195–205.

Tugiman ES, Yusoff MZM, Hassan MA, Abd Samad MY, Farid MAA, Shirai Y (2024) Assessing the efficacy of utilizing biochar derived from oil palm biomass as a planting medium for promoting the growth and development of oil palm seedlings. Biocatalysis and Agricultural Biotechnology, 58, 103203. 10.1016/j.bcab.2024.103203

Uselman SM, Qualls RG, Thomas RB (2000). Effects of increased atmospheric CO_2_, temperature, and soil N availability on root exudation of dissolved organic carbon by a N-fixing tree (*Robinia pseudoacacia* L.). Plant and Soil, 222, 191–202. 10.1023/A:1004705416108

van de Voorde TF, Bezemer TM, Van Groenigen JW, Jeffery S, Mommer L (2014) Soil biochar amendment in a nature restoration area: effects on plant productivity and community composition. Ecological Applications, 24, 1167–1177. 10.1890/13-0578.1

Viechtbauer W (2010) Conducting meta-analyses in R with the metafor package. Journal of Statistical Software, 36, 1–48. 10.18637/jss.v036.i03

Wan Mohamed WNA, Liu CL, Abdullah R, & Osman N (2021). Influence of organic amendments on the growth and physiology of *Melastoma malabathricum*. Transactions of the Malaysian Society of Plant Physiology, 28, 79–85.

Wang B, Lehmann J, Hanley K, Hestrin R, Enders A (2015). Adsorption and desorption of ammonium by maple wood biochar as a function of oxidation and pH. Chemosphere, 138, 120–126. 10.1016/j.chemosphere.2015.05.062

Wang X, Zheng WL, Ma X, Yu FH, Li MH (2023) Biochar aggravates the negative effect of drought duration on the growth and physiological dynamics of *Pinus massoniana*. Frontiers in Ecology and Evolution, 11, 1166538. 10.3389/fevo.2023.1166538

Williams JM, Thomas SC (2023) Effects of high-carbon wood ash biochar on volunteer vegetation establishment and community composition on metal mine tailings. Restoration Ecology, 31, e13861. 10.1111/rec.13861

Wijaya BA, Riniarti M, Prasetia H, Hidayat W, Niswati A, Hasanudin U, Banuwa IS (2021) Interaksi perlakuan dosis dan suhu pirolisis pembuatan biochar kayu meranti (*Shorea* spp.) mempengaruhi kecepatan tumbuh sengon (Paraserianthes moluccana). ULIN: Jurnal Hutan Tropis, 5, 78–89. (in Indonesian). 10.32522/ujht.v5i2.5782

Wijaya BA, Hidayat W, Riniarti M, Prasetia H, Niswati A, Hasanudin U, Banuwa IS, Kim S, Lee S, Yoo J (2022) Meranti (*Shorea* sp.) biochar application method on the growth of sengon (Falcataria moluccana) as a solution of phosphorus crisis. Energies, 15, 2110. 10.3390/en15062110

Xiao Y, Wang L, Zhao Z, Che Y (2020) Biochar shifts biomass and element allocation of legume-grass mixtures in Cd-contaminated soils. Environmental Science and Pollution, 27, 10835–10845. 10.1007/s11356-019-07357-3

Yaebiyo G, Birhane E, Tadesse T, Kiros S, Hadgu KM, Egziabher YG, Habtu S (2023) Using biochar and deficit irrigation enhanced the growth of commercial agroforestry woody species seedlings in drylands (a case study in Saz, northern Ethiopia). Agroforestry Systems, 98, 61–79. 10.1007/s10457-023-00891-7

Ye L, Camps-Arbestain M, Shen Q, Lehmann J, Singh B, Sabir M (2020) Biochar effects on crop yields with and without fertilizer: a meta-analysis of field studies using separate controls. Soil Use and Management, 36, 2–18. 10.1111/sum.12546

Yousaf MTB, Nawaz MF, Zia ur Rehman M, Gul S, Yasin G, Rizwan M, Ali S (2021) Effect of three different types of biochars on eco-physiological response of important agroforestry tree species under salt stress. International Journal of Phytoremediation, 23, 1412–1422. 10.1080/15226514.2021.1901849

Yousaf MTB, Nawaz MF, Zia M, Rasul F& Tanvir MA (2021) Ecophysiological response of early stage *Eucalyptus camaldulensis* to biochar and other organic amendments under salt stress. Pakistan Journal fo Agricultural Science, 58, 999–1006. 10.21162/PAKJAS/21.1012

Zou Z, Mi W, Li X, Hu Q, Zhang L, Zhang L, Fu J, Li Z, Han W, Yan P (2023) Biochar application method influences root growth of tea (*Camellia sinensis* L.) by altering soil biochemical properties. Scientia Horticulturae, 315, 111960. 10.1016/j.scienta.2023.111960

